# HIV-1 prime-boost vaccination shapes distinct clonal trajectories and memory precursor states of Env- and Gag-specific T cells

**DOI:** 10.64898/2026.09.23.752815

**Authors:** Guoyue Xu, Joshua X. Yee, Saskia M. Ilcisin, Saransh N. Kaul, Koshlan Mayer-Blackwell, Ernest Moelhman, David R. Glass, Anastasia A. Minervina, Shelly T. Karuna, Magdalena E. Sobieszczyk, Matthew R. Hart, Paul G. Thomas, Peter J. Skene, Stephen C. De Rosa, M. Juliana McElrath, Andrew Fiore-Gartland, Evan W. Newell

## Abstract

Despite decades of HIV-1 vaccine development, the clonal and cellular determinants of durable vaccine-induced T cell memory remain incompletely understood. Here, we combined antigen-specific T cell receptor (TCR) identification, longitudinal TCR sequencing, and single-cell multi-omics to characterize Env- and Gag-specific memory precursor T cells elicited by the HIV Vaccine Trials Network (HVTN) 505 DNA prime–recombinant adenovirus serotype 5 (rAd5) boost (DNA/rAd5) vaccine regimen. We developed a generalizable high-throughput approach to identify HIV-1 Env- and Gag-specific TCRs and found distinct patterns of clonal expansion, persistence, and contribution to memory between Env- and Gag-specific CD8⁺ T cell responses. The DNA prime and rAd5 boost differentially shaped these repertoires, with rAd5-induced clones contributing proportionally more to the Gag-specific than to the Env-specific memory precursor compartment. Single-cell immune profiling further revealed distinct memory precursor states, with Env-specific responses enriched for *GZMB^+^PRF1^+^* cytotoxic effector-memory (EM) CD8⁺ T cells and Gag-specific responses containing a larger cycling/proliferative population. Together, these findings demonstrate that heterologous DNA/rAd5 vaccination generates antigen-specific CD8⁺ T cell memory with distinct clonal trajectories and cellular programs, providing new insights into how vaccine platform and antigen-specificity shape the durability and functional properties of HIV-1 specific cellular immunity.

## INTRODUCTION

Despite significant progress in HIV-1 vaccine research, a licensed vaccine has remained elusive. A major challenge is generating durable protective immunity against a genetically diverse and rapidly evolving virus (*1*). While current HIV-1 vaccine efforts have focused largely on broadly neutralizing antibodies, evidence suggests that strong and durable T cell mediated responses may also contribute to protection (*1*, *2*). Prior HIV-1 vaccine efficacy trials provide a unique opportunity to apply high-resolution sequencing technologies to define vaccine-induced cellular immune responses and identify biomarkers that may inform future vaccine design.

Traditional immunological methods used to assess vaccine immunogenicity do not resolve clonal-level dynamics of the cellular immune response (*3*). TCR sequencing, by contrast, enables tracking of vaccine-induced T cell clones and has revealed clonal determinants of protective immunity following yellow fever and SARS-CoV-2 vaccination (*4–6*). In HIV-1, cellular immunity is well established as an important component of viral control, particularly the contribution of Gag-specific CD8⁺ T cells (*7*, *8*). Efforts to characterize HIV-1-specific T cell repertoires have identified TCRs recognizing immunodominant epitopes such as the p24 Gag epitope (*9*, *10*). However, most TCR studies in HIV-1 have focused on individual epitopes, specific HLA alleles, or people living with HIV-1 (*10–13*). Thus, the broader antigen-specific T cell repertoire elicited by HIV-1 vaccination, particularly its clonal persistence and relationship to memory formation, remains incompletely characterized.

Durable immune memory is a central goal of vaccination. Antigen-specific memory T cells rapidly expand and acquire effector function following antigen re-exposure (*14*, *15*). Both the effector and memory T cell compartments exhibit substantial phenotypic and functional heterogeneity that influences immune protection (*16*). The transition from activated effector T cells to long-lived memory is dynamic, and only a small subset of effector clonotypes successfully become durable memory precursors (*15*, *17*). For HIV-1 vaccines, it therefore remains unclear which vaccine-induced T cell populations persist and contribute to long-term memory.

Although the RV144 trial remains the only HIV-1 vaccine efficacy trial to demonstrate modest protection, evidence for the role of cytotoxic CD8⁺ T cells in HIV-1 control motivated the development of multiple T cell-focused vaccine strategies (*18*, *19*). Immune correlate analyses from the rAd5-based Merck 023/ HVTN 502 and HVTN 505 trials identified associations between cytotoxic CD8⁺ T cells responses and reduced viral load or HIV-1-acquisition (*20–22*). Subsequent analyses of HVTN 505 identified the magnitude and polyfunctionality of Env-specific CD8⁺ T cell responses as immune correlates associated with lower rates of HIV-1-infection (*20*, *23*, *24*). However, the durability, specificity, and cellular phenotypes of HIV-1-specific T cells induced by the DNA/rAd5 vaccine remain poorly characterized. We therefore sought to compare the clonal dynamics, cellular phenotypes, and molecular programs of vaccine-induced Env- and Gag-specific T cell responses.

In this study, we developed a high-throughput approach to identify vaccine-induced antigen-specific TCRs in HVTN 505 vaccine recipients. Mapping these TCRs against longitudinal TCR and single-cell multi-omics data revealed that prime-boost DNA and rAd5 vaccination modalities differentially shaped the expansion, persistence and cellular states of Env- and Gag-specific CD8⁺ T cells. These cells exhibited distinct cellular states and clonal trajectories, with Env- specific responses enriched for *GZMB^+^PRF1^+^* cytotoxic EM features and Gag-specific responses showing greater representation of cycling/proliferative cells. Together, these findings define how HIV-1 vaccine-induced T cell clones evolve over time and link clonal trajectories with cellular states associated with durable immune memory.

## RESULTS

### Generation and validation of high-confidence HIV-1 Env- and Gag-specific TCR repertoires

HVTN 505 was a phase 2b efficacy trial that evaluated a heterologous multi-gene, multi-clade DNA/rAd5 HIV-1 prime-boost vaccine regimen in the United States (*20*, *21*). The DNA prime consisted of HIV-1 Gag, Pol, and Nef from clade B and Env from clades A, B, and C. Participants subsequently received a rAd5 boost at month 6 (M6) expressing an HIV-1 clade B Gag-Pol fusion protein and Env glycoproteins from clades A, B, and C (Fig. 1A) (*21*). Although the vaccine did not demonstrate efficacy, subsequent analyses found that higher-magnitude and more polyfunctional Env-specific CD8⁺ T cell responses were associated with reduced risk of HIV- 1 acquisition (*20*). For this study, we selected 16 vaccine recipients who remained HIV-1-negative and demonstrated vaccine-induced immune responses to at least one Env or Gag T cell condition by intracellular cytokine staining (ICS) (table S1). Individual participants did not necessarily exhibit detectable responses across all antigen and cell-type combinations.

**Fig. 1.**
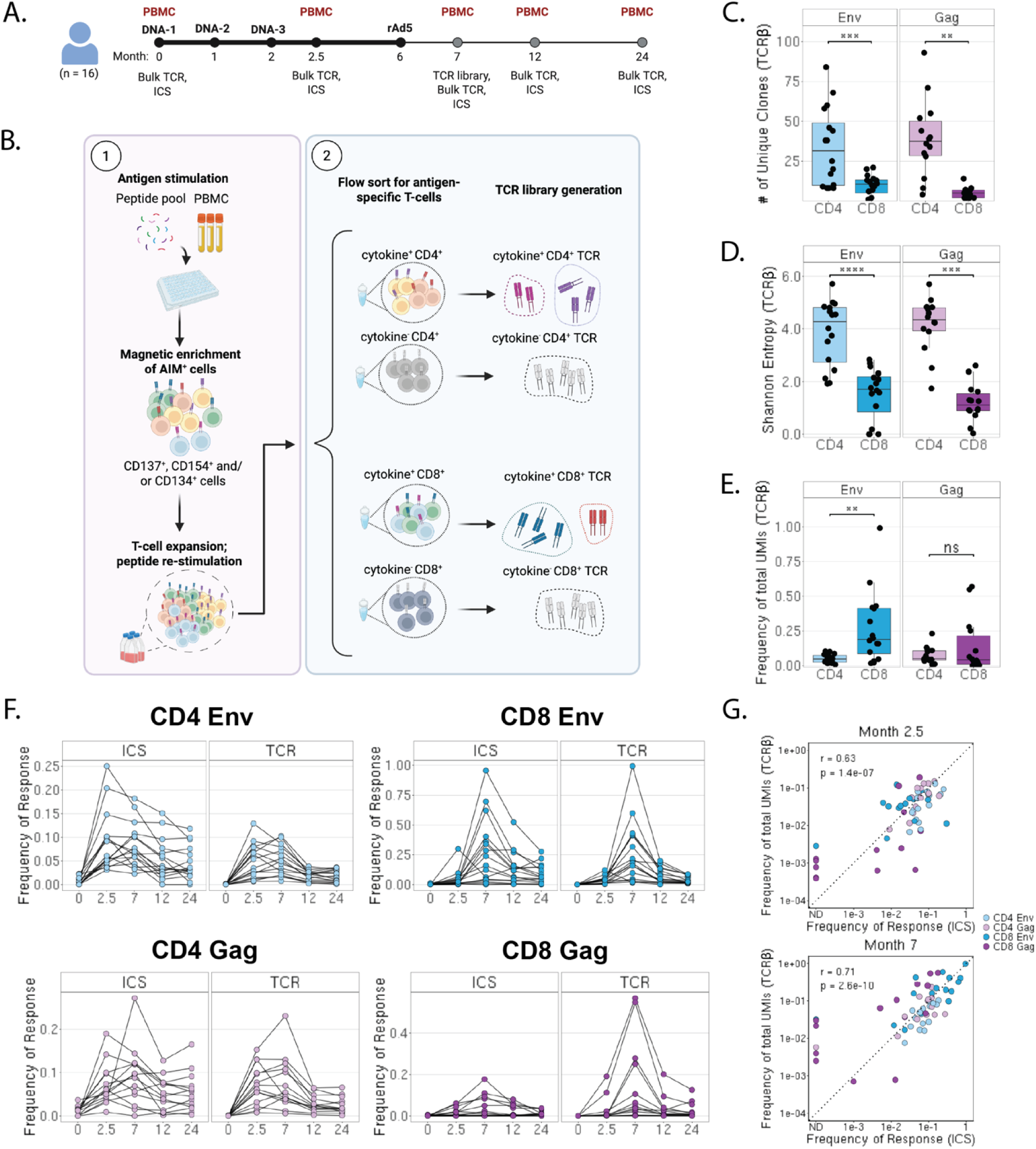
Isolation and identification of vaccine-specific HIV-1 Env- and Gag-specific TCRs. **A)** Schematic of the HVTN 505 vaccination regimen, sample collection time points, and experimental design **B)** Overview of the experimental strategy used to isolate and generate antigen-specific TCR libraries **C)** Number of unique vaccine-specific TCRβ clones identified per participant at M7, shown for Env- and Gag-specific CD4⁺ and CD8⁺ T cells **D)** Shannon entropy of vaccine-specific TCRβ clones at M7, calculated from the relative frequencies of individual TCRβ clones per participant **E)** Magnitude of Env- and Gag-specific CD4⁺ and CD8⁺ T cell responses at M7, measured as the frequency of vaccine-specific TCRβ UMI counts within the total TCRβ repertoire for each participant **F)** Longitudinal vaccine-specific T cell responses measured by TCR sequencing and ICS for each participant. Lines connect measurements from the same participant. For TCR sequencing, response magnitude represents the frequency of vaccine-specific TCRβ UMI counts within the total TCRβ repertoire; for ICS, response represents the frequency of cytokine-responsive T cells **G)** Correlation between vaccine-specific T cell responses measured by TCR sequencing and ICS across participants at M2.5 and M7 for Env- and Gag-specific CD4⁺ and CD8⁺ T cells. Each point represents an individual participant. Spearman rank correlation coefficients and *P* values are shown. Data are plotted on logarithmic scales Statistical comparisons in (C)–(E) were performed using the paired Wilcoxon signed-rank test. \**P* < 0.05; \*\**P* < 0.01; \*\*\**P* < 0.001; \*\*\*\**P* < 0.0001; ns, not significant; ND, not detectable.

We profiled vaccine-induced Env- and Gag- specific TCR repertoires from cryopreserved PBMC samples at M7 (Fig. 1A). Because antigen-specific TCRs cannot be readily identified from sequence alone, we developed a two-stage approach to enrich, expand and identify rare antigen-specific T cells (Fig. 1B). We stimulated PBMCs using vaccine-matched Env or Gag peptide pools, followed by magnetic enrichment of T cells expressing any combination of activation-induced markers (AIM) and *in vitro* expansion using a rapid-expansion protocol (*25*, *26*). Expanded AIM^+^ cells were re-stimulated with the same peptide pools and antigen-specific T cells (cytokine^+^) were identified using a three-cytokine Cytokine Secretion Assay (CSA) and sorted for bulk TCRα/β chain sequencing based on their cytokine secretion (fig. S1A)(see Supplemental Materials and Methods)(*27*). Expanded T cells and cytokine non-secreting cells were sequenced in parallel as controls. Unstimulated longitudinal PBMC samples collected before vaccination (M0), post-DNA prime (M2.5), post-vaccination (M7) and two durability time-points (M12; M24) were also sequenced (Fig. 1A).

To identify high-confidence vaccine-specific clones, we selected TCRs showing sequential enrichment from the longitudinal PBMC repertoire through the subsequent expansion and cytokine^+^ populations, together with higher frequency in the post-expansion cytokine^+^ vs. the cytokine^-^ population (fig. S1B) (see Supplemental Materials and Methods). We further required all cytokine^+^ clones to have fewer than 2 Unique Molecular Identifier (UMI) counts in the corresponding M7 PBMC sample. Mapping these TCRs to the longitudinal repertoire identified 1439 unique TCRα chains and 1680 unique TCRβ chains corresponding to HIV-1 Env- and Gag-specific CD4⁺ and CD8⁺^.^ T cells (table S2 and S3).

The vaccine expanded significantly more unique Env- and Gag- specific CD4⁺ than CD8⁺ T cell clones per participant at M7 (Env, *P* = 0.0007; Gag, *P* = 0.001; Fig. 1C), resulting in greater clonal diversity among CD4⁺ T cells (Env, *P* = 3.05x10^-5^; Gag, *P* = 1.22x10^-4^; Fig. 1D). Despite the greater clonal diversity of CD4⁺ T cells, the total frequency of Env- specific CD8⁺ T cells at M7 was higher than that of Env-specific CD4⁺ T cells (Env, *P* = 0.001, Fig. 1E), consistent with a more oligoclonal CD8⁺ T cell response dominated by expanded clones (fig. S2A).

We next calculated the cumulative frequency of Env- and Gag-specific TCR clones at each time point and compared TCR-derived response frequencies with ICS measurements to validate the antigen-specific TCR repertoire (Fig. 1F, and table S4)(*28*). TCR- and ICS- derived response magnitudes showed strong concordance across participants, time points and T cell subsets (Fig. 1G and fig. S2B). However, TCR-sequencing identified substantially greater Gag-specific CD8⁺ T cell frequencies than ICS in a subset of participants, suggesting that TCR-based measurements can capture antigen-specific clones that could not be detected by cytokine-based assays. Together, these results establish a high-confidence repertoire of vaccine-induced HIV-1 Env- and Gag-specific TCRs and demonstrate that TCR-based measurements can recapitulate conventional immunogenicity measurements while providing clonotypic resolution.

### DNA-prime and rAd5-boost differentially shape Env- and Gag-specific CD8⁺ T cell expansion and clonal composition

Evaluating the longitudinal effects of HVTN 505 vaccination, we found that CD4⁺ and CD8⁺ T cell responses exhibited distinct expansion dynamics following prime and boost immunization (Fig. 2A). Most participants’ CD8⁺ T cell responses peaked following the rAd5 boost, whereas CD4⁺ T cell responses showed a more even distribution of peak responses (Fig. 2B; *P* = 0.01, quantified in fig. S3A).

**Fig. 2.**
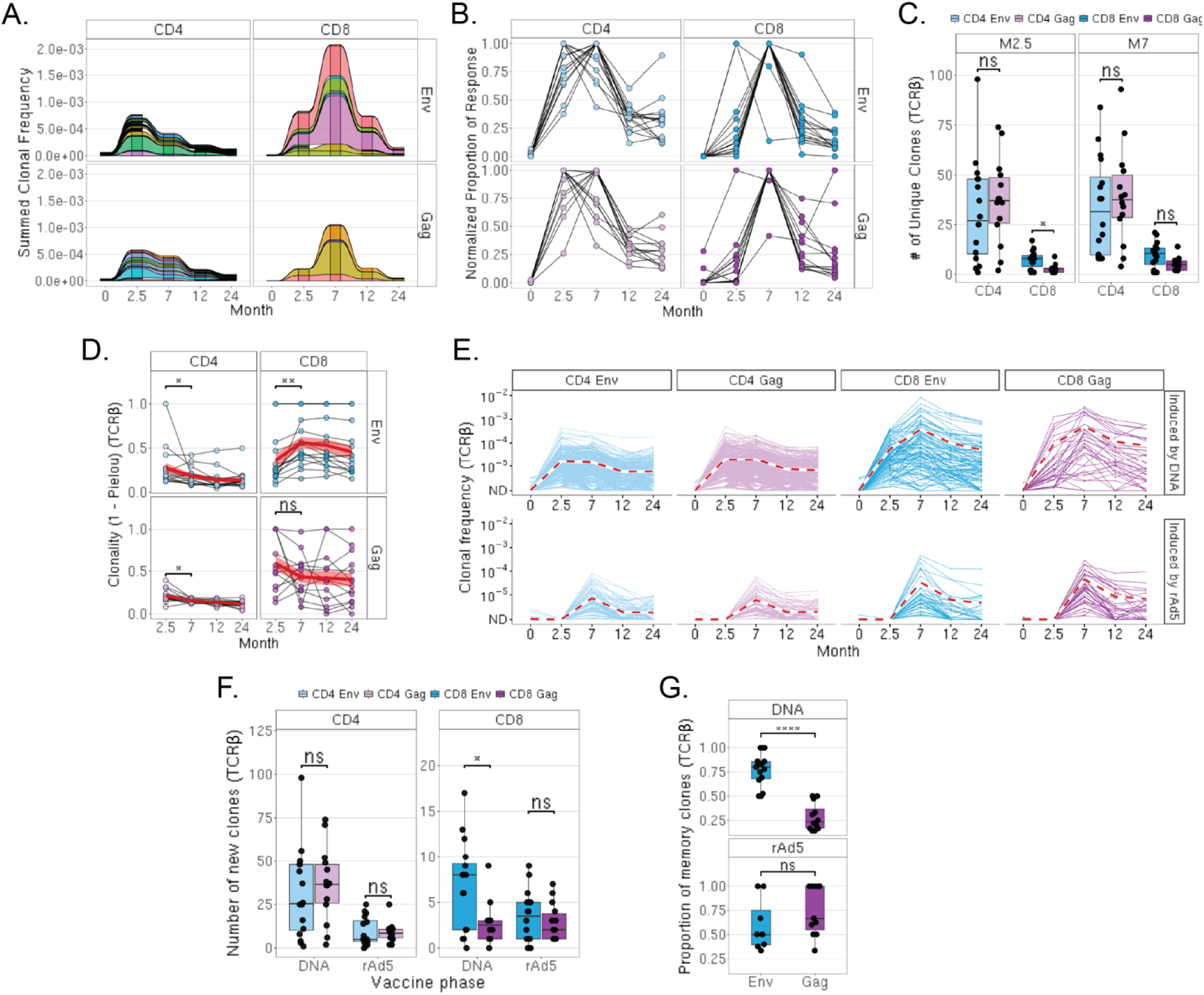
DNA prime and rAd5 boost differentially shape the Env- and Gag-specific T cell repertoire **A)** Longitudinal dynamics of the top 10 clones ranked by frequency at each time point for vaccine-specific CD4⁺ and CD8⁺ T cells responding to Env or Gag. Each alluvial flow represents an individual TCRβ clone, with clone frequency shown across time points **B)** Longitudinal TCRβ response magnitude normalized to the peak response for each participant and cell type–antigen combination. Individual lines connect measurements from the same participant **C)** Number of unique TCRβ clones in Env- and Gag-specific CD4⁺ and CD8⁺ T-cell responses at M2.5 and M7. Boxplots show the distribution across participants and individual points represent participants. Statistical comparisons between Env- and Gag-specific responses were performed using an unpaired Wilcoxon rank-sum test **D)** Longitudinal TCRβ repertoire clonality, calculated as 1 − Pielou’s evenness. Individual points represent participants and lines connect paired measurements. Red lines and shaded areas indicate the mean ± standard error. Paired comparisons between M2.5 and M7 were performed using the Wilcoxon signed-rank test, with *P-*values adjusted using the BH method **E)** Longitudinal trajectories of vaccine-specific TCRβ clones classified according to the time of initial expansion. Clones were classified as DNA-induced or rAd5-induced based on the time-point they were first detected. Thin lines represent individual clones and dash red lines indicate the mean trajectory for each expansion group **F)** Number of newly expanded TCRβ clones induced following DNA prime or rAd5 boost, shown separately for CD4⁺ and CD8⁺ T cells. Each point represents a participant. Paired comparisons between Env- and Gag-specific responses were performed using the Wilcoxon signed-rank test, with *P*-values adjusted using the BH method **G)** Proportion of memory TCRβ clones among clones initially induced by DNA prime or rAd5 boost. Analyses are shown separately for Env- and Gag-specific CD8⁺ T cells. Boxplots show the distribution across participants and individual points represent participants. Comparisons between Env- and Gag-specific responses within each vaccine phase were performed using the Wilcoxon rank-sum test, with *P*-values adjusted using the BH method *\*P* < 0.05; \*\**P* < 0.01; \*\*\**P* < 0.001; \*\*\*\**P* < 0.0001; ns, not significant; ND, not detectable.

Prime-boost vaccination also differentially shaped the clonal composition of antigen-specific T cell responses. Although the number of detected Env- and Gag- specific clones increased across T cell populations following DNA prime, more Env-specific CD8⁺ T cell clones were detected (Fig. 2C and fig. S3B). Following rAd5 boost, clonality decreased across CD4^+^ (Env and Gag) and CD8^+^ (Gag) T cell populations, consistent with additional exposure leading to repertoire diversification; however, the Env-specific CD8⁺ T cell repertoire became more clonal (Fig. 2D), suggesting preferential expansion of select clones recognizing this antigen.

To further define how vaccination shaped the vaccine-induced T cell repertoire, we categorized clones by the timepoint at which they were first detected and tracked their longitudinal clonal trajectories (Fig. 2E). Following DNA prime, significantly fewer new Gag-specific CD8⁺ T cell clones were detected compared with Env-specific clones (Fig. 2F). In contrast, rAd5-boost recruited comparable numbers of newly detected Env- and Gag-specific CD8⁺ T cell clones. We next defined clones detected at M24 as the memory compartment and found that DNA prime-induced Gag-specific clones accounted for a smaller proportion of the memory CD8⁺ T cell repertoire, whereas Env-specific memory was composed predominantly of clones recruited following the DNA prime (Fig. 2G). DNA prime-induced Gag-specific CD8⁺ T cell clones showed a trend toward faster contraction than Env-specific clones, which may contribute to their reduced representation in long-term memory (fig. S3C). Together, these findings demonstrate that DNA prime preferentially established Env-specific CD8⁺ T cell clones that contributed substantially to long-term memory, whereas rAd5 boost recruited new clones that contributed proportionally more to the Gag-specific memory compartment and expanded established Env-specific clones.

### Vaccine-specific clonotypes can be identified through unbiased trajectory analyses

Although our experimental approach identified clade B Env- and Gag-specific TCRs, we sought to define the broader vaccine-induced repertoire using an unbiased longitudinal approach. We compared clonal frequencies at M2.5 and M7 with baseline (M0), classifying clones as vaccine-expanded based on a log2 fold-change >2 and Benjamini-Hochberg (BH)-adjusted Fisher’s exact test *P* < 0.05 (Fig. 3A and fig. S4A)(*29*, *30*). This approach identified highly expanded clones but failed to capture many low-frequency Env and Gag-specific clones identified experimentally by our two-stage method (Fig. 3A). Conversely, statistical criteria identified dynamic clones that were already abundant at baseline, potentially reflecting pre-existing or cross-reactive T cells rather than newly induced vaccine-specific responses (*31–33*). Consequently, statistically enriched clones only partially overlapped with experimentally defined Env- and Gag-specific TCRs (fig. S4B), demonstrating limited sensitivity and specificity for identifying the antigen-specific repertoire.

**Fig. 3.**
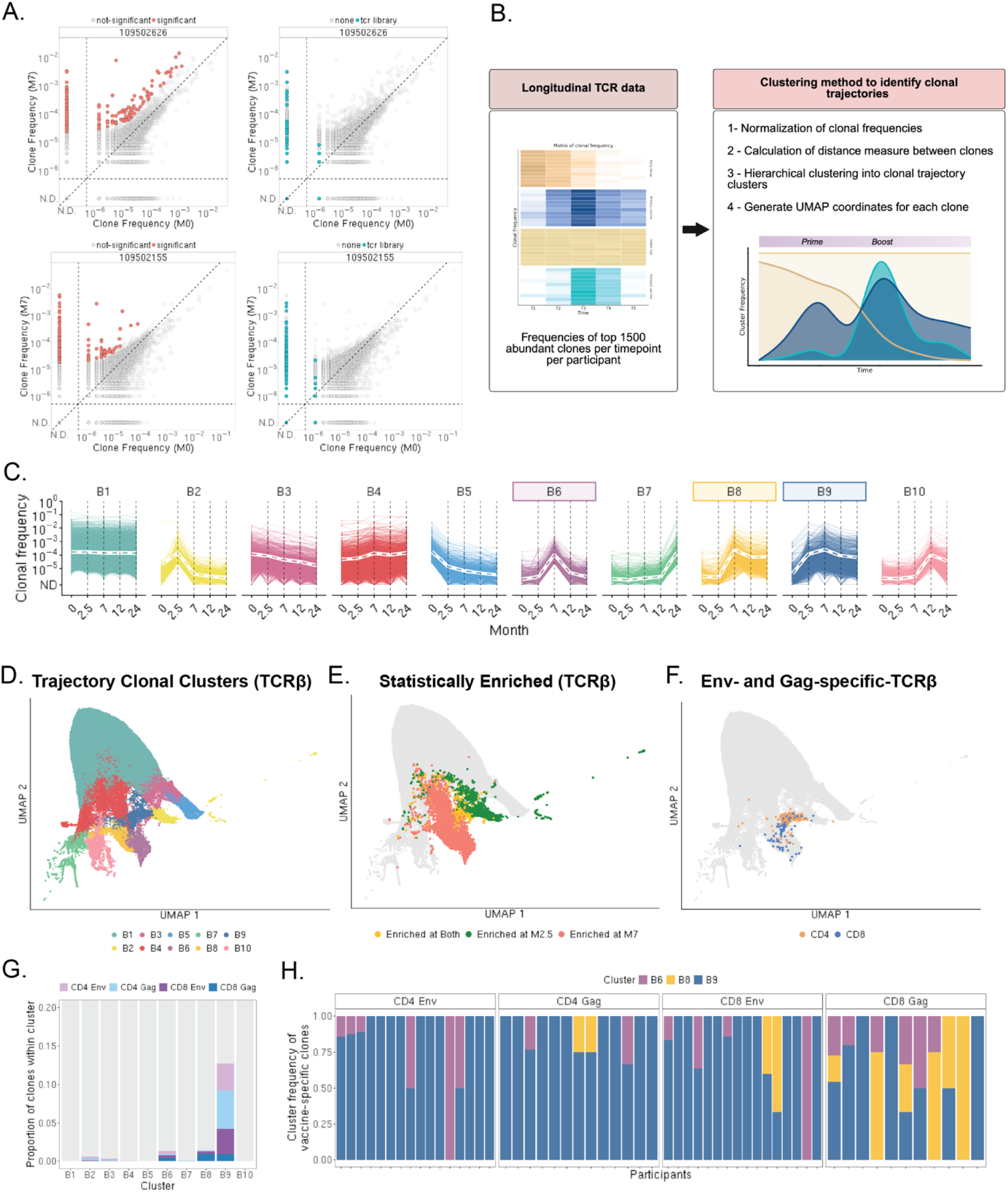
Unbiased trajectory analysis identifies durable vaccine-induced clonal kinetics **A)** Biaxial plots comparing TCRβ clonal frequencies at M0 and M7 for two representative participants. *C*lones that meet the statistical cut-off criteria of log2 fold-change (FC) >2 and FDR-adjusted *P-*value < 0.05 by Fisher’s exact test are shown in coral (*left)* and experimentally identified HIV-1 Env- and Gag-specific TCRβ clones are shown in teal (*right*) **B)** Schematic of the longitudinal trajectory analysis used to identify distinct clonal kinetic patterns (see Materials and Methods for details) **C)** Longitudinal TCRβ clonal trajectories grouped into ten clusters representing distinct patterns of expansion and persistence. Colored boxes indicate trajectory clusters significantly enriched for Env- or Gag-specific TCRβs **D)** UMAP representation of the longitudinal TCRβ clonal trajectories shown in (C), colored according to trajectory cluster **E)** UMAP colored by clones identified as statistically enriched at M2.5 (green), M7 (coral), or both time points (yellow) **F)** UMAP colored by experimentally identified vaccine-specific CD4⁺ (orange) and CD8⁺ (blue) TCRβs from the antigen-specific TCR libraries **G)** Stacked bar plot showing the number of antigen-specific cells within each trajectory cluster, stratified by antigen and T-cell subset **H)** Stacked bar plots showing the proportion of vaccine-specific TCRβs assigned to each trajectory cluster for each participant ND, non-detectable

To overcome these limitations, we applied an unbiased trajectory-based clustering framework to characterize longitudinal clonal kinetics (Fig. 3B)(see Materials and Methods)(*6*, *30*, *34*). Hierarchical clustering of longitudinal TCRβ frequencies resolved 10 distinct trajectory clusters (B1-B10) representing stable (B1, B3, and B4), transient (B5, B7, and B10), and hypothesized-vaccine-responsive patterns of clonal behavior (B2, B6, B8, and B9) (Fig. 3C) . Clones within each cluster localized to corresponding regions of the UMAP, supporting the trajectory classification (Fig. 3D). Statistically enriched clones were distributed across multiple clusters, with many localized to B4, a cluster characterized by persistent abundance throughout the study period (Fig. 3, C and E). We next applied the same trajectory analysis independently to the TCRα repertoires and observed similar patterns of clonal behavior (fig. S4, C-D).

We next asked whether experimentally validated vaccine-specific TCRs were preferentially associated with specific trajectory clusters. Env- and Gag-specific TCRs were significantly enriched within clusters B6, B8, and B9 and localized to corresponding regions of the trajectory UMAP (Fig. 3, F and G; table S5). These clusters exhibited kinetics consistent with vaccine-induced expansion, with clones increasing from low baseline frequencies following either DNA prime or rAd5 boost (Fig. 3C). Clusters B8 and B9 contained the majority of long-lived vaccine-induced clones detectable through M24, whereas B6 showed short-lived kinetics with rapid post-expansion contraction. Because experimentally defined TCR libraries were generated at M7, transient clones induced exclusively by DNA prime may not have been captured and could contribute to cluster B2.

Consistent with our finding that Gag-specific CD8⁺ T cell memory includes clones recruited by both DNA prime and rAd5 boost (Fig. 2H), Gag-specific CD8⁺ TCRs were more frequently associated with the rAd5-responsive memory trajectory B8 than other vaccine-specific populations (Fig. 3, G and H). Together, these findings identify B6, B8, and B9 as the principal vaccine-responsive trajectories, with B8 and B9 representing the dominant long-lived memory trajectories following DNA/rAd5 vaccination. Clones within these trajectories that were not experimentally assigned to clade B Env or Gag may represent T cells targeting other vaccine-encoded antigens with similar longitudinal kinetics.

### Prime-boost DNA/rAd5 HIV-1-vaccination induces cytotoxic and proliferative CD8⁺ T cell populations with distinct clonal trajectories

We used the newly developed Reverse-transcription Enabled-FLEX (REFLEX) platform to define the cellular and molecular features associated with distinct clonal expansion and persistence kinetics following heterologous DNA/rAd5 HIV-1 vaccination (*38*). REFLEX enables simultaneous profiling of gene expression, surface protein expression using antibody-derived tags (ADTs), and paired-chain TCR sequences of fixed cells. We profiled total CD3^+^ T cells isolated from M7 PBMC samples from 11 of the 16 original study participants (table S1). After quality control and removal of non-T cell populations, 294,583 T cells were retained for downstream analyses, including 244,318 cells with TCR information.

To integrate the single-cell and longitudinal bulk TCR datasets, we re-performed clonal trajectory clustering on the top 1,500 clones identified in Fig. 3, restricting the analysis to clones detected in the REFLEX dataset (fig. S5A). These analyses identified five vaccine-induced trajectory clusters, including SC1, SC10, and SC12, which had previously been grouped within trajectory cluster B9 in the bulk longitudinal clonal analysis (Fig. 4A and fig. S5A). Consistent with our longitudinal analyses and previous reports describing distinct expansion kinetics of vaccine-induced CD4⁺ and CD8⁺ T cells, clones were differentially distributed across trajectory clusters: SC1 and SC12 were enriched for CD4⁺ T cells, SC7 and SC10 for CD8⁺ T cells, and SC9 contained comparable proportions of both populations (fig. S5B) (*39*, *40*). To define the cellular states associated with these trajectories, we performed weighted nearest neighbor (WNN) analysis on CD4⁺ and CD8⁺ T cells separately using integrated transcriptomic and surface protein data (*41*), identifying nine CD4⁺ T cell clusters and twelve CD8⁺ T cell phenotypic clusters (Fig. 4B and fig. S6)(*17*). Participant-matched TCR sequences were then used to map vaccine-specific clones from the trajectory clusters onto the single-cell dataset (Fig. 4C), enabling direct integration of longitudinal clonal behavior with cellular phenotype.

**Fig. 4.**
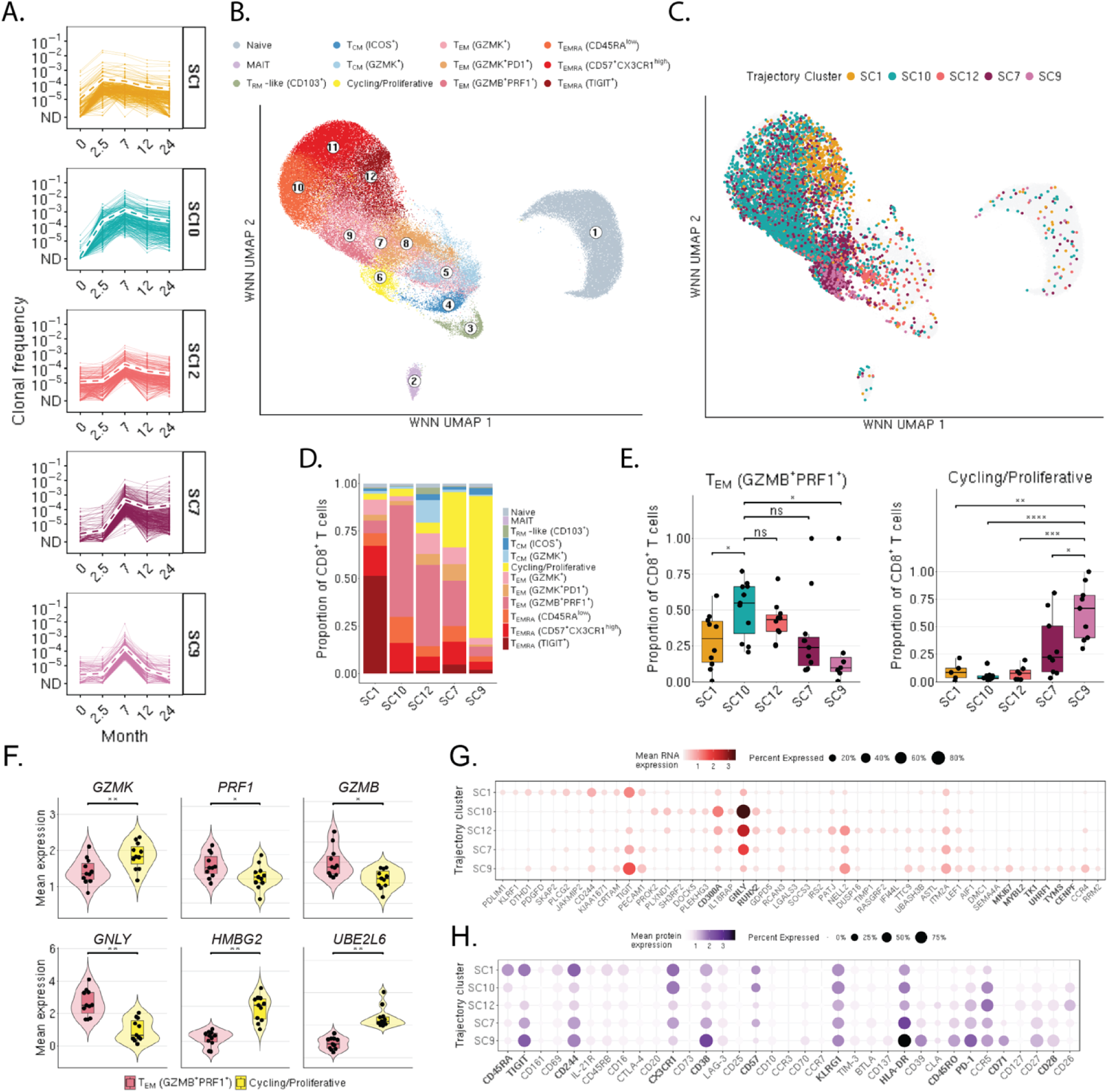
Distinct clonal trajectories are associated with cytotoxic and proliferative CD8⁺ T cells following DNA/rAd5 HIV vaccination A) Vaccine-specific trajectory clusters identified by hierarchical clustering following integration of single-cell TCR clones with the longitudinal vaccine-specific TCR datasets B) CD8⁺ T cell UMAP colored by annotated T cell states C) CD8⁺ T cell UMAP colored by longitudinal trajectory clusters from (A) D) Stacked bar plot showing the proportion of CD8⁺ T cell states within each trajectory cluster E) Participant-level proportion of cells within each trajectory cluster belonging to the indicated CD8⁺ T cell state. Differences between trajectory clusters were assessed using pairwise Wilcoxon rank-sum tests with BH correction for multiple comparisons F) Violin plots showing expression of selected genes associated with *GZMB^+^PRF1^+^* T_EM_ and cycling/proliferative CD8⁺ T cell states. Gene expression values were averaged across cells for each participant within each T cell state. Differences between states were assessed using Wilcoxon signed-rank tests with BH correction for multiple comparisons G) RNA expression profiles of the top significantly enriched genes within each single-cell trajectory cluster. Cluster-enriched genes were identified using one-versus-all differential expression analysis and visualized as dot plots. Dot size represents the percentage of cells expressing each gene, and color indicates mean RNA expression within each trajectory cluster H) Protein expression profiles of the top significantly enriched ADT markers within each single-cell trajectory cluster. Differentially abundant surface proteins were identified using one-versus-all analysis and visualized as dot plots. Dot size represents the percentage of cells positive for each protein, and color indicates mean protein expression within each trajectory cluster *\* P* < 0.05; ** *P* < 0.01; *** *P* < 0.001; **** *P* < 0.0001; ns, not significant; ND, non-detectable

Trajectory and phenotypic associations were most pronounced within the CD8⁺ T cell compartment in contrast to the CD4⁺ T cell compartment (fig. S7). Given previous HVTN 505 studies linking Env-specific CD8⁺ T cell responses with reduced HIV-1 acquisition risk, together with the established importance of Gag-specific CD8⁺ T cells in HIV control, we focused subsequent analyses on the CD8⁺ T cell compartment (*7*, *8*, *20*). The response was dominated by two major phenotypic states: differentiated cytotoxic *GZMB^+^PRF1^+^*T_EM_ and activated cycling/proliferative cells (Fig. 4D and fig. S8A). Except for SC1, whose enrichment for *TIGIT^+^* T effector memory cells re-expressing CD45RA (T_EMRA_) cells was driven by a single participant (fig. S8A), DNA prime-induced trajectories SC1 and SC10 and rAd5-induced SC12 were enriched for differentiated cytotoxic *GZMB^+^PRF1^+^* T_EM_ cells, whereas rAd5-induced SC7 and SC9 were enriched for activated cycling/proliferative cells (Fig. 4E). Notably, these phenotypic states reflected distinct clonal histories: SC10 was composed predominantly of clones primed by DNA vaccination and subsequently expanded following rAd5 boost, whereas SC9 was largely composed of rAd5-induced clones. The *GZMB^+^PRF1^+^*T_EM_ population expressed canonical cytotoxic genes including *GZMB, GZMH, GNLY, PRF1, NKG7,* and *GZMK*, whereas cycling/proliferative cells retained effector-associated genes such as *PRF1*, *NKG7*, *CCL5*, and *GZMK* but exhibited increased *HMGB2* and *UBE2L6* expression and reduced expression of *GZMB*, *GNLY,* and *KLRD1* (Fig. 4F and fig. S6C)(*15*, *42*). The activated phenotype of cycling/proliferative cells was further supported by increased surface expression of CD38, HLA-DR, CD39, PD-1, and TIGIT (Fig. 4F and fig. S6, A and B)(*6*, *42–44*).

Differentially expressed genes (DEGs) and differentially expressed proteins (DEPs) analyses further distinguished these trajectory-associated phenotypes. SC10 exhibited the strongest cytotoxic transcriptional program, with elevated expression of *GNLY, CD300A*, and *RUNX2*, genes associated with cytotoxic function, immune regulation, and memory differentiation (Fig. 4G) (*45–47*). In contrast, SC9 showed a proliferative program characterized by increased expression of cell-cycle and DNA replication genes including *MKI67, TYMS, CENPF, UHRF1, MYBL2,* and *TK1* (*41*, *48*). SC7 and SC12 displayed intermediate transcriptional programs, combining cytotoxic features with activation- and proliferation-associated genes. DEPs analyses similarly distinguished SC10 from the rAd5-associated trajectories, with increased expression of CX3CR1, CD57, and CD45RA and reduced expression of CD28, CD45RO, and TIGIT (Fig. 4H). In contrast, SC7, SC9, and SC12 showed increased expression of activation-associated proteins including HLA-DR, CD38, CD71, and CD244, together with higher levels of PD-1, TIGIT, and KLRG1 (Fig. 4H) (*49*, *50*). Together, these analyses support SC10 as a differentiated cytotoxic memory population with features of durable memory, whereas SC9 represents a highly activated proliferative state, and SC7 and SC12 displayed intermediate phenotypes.

Collectively, these findings identify distinct vaccine-induced CD8⁺ T-cell populations associated with divergent clonal trajectories, phenotypic states, and persistence kinetics. Despite substantial contribution of vaccine-expanded clones to the durable memory compartment, the highly proliferative SC9 population displayed comparatively limited durability.

### Distinct clonal trajectories underlie divergent states of Env- and Gag-specific CD8⁺ memory precursor T cells

We next investigated whether vaccine-induced Env- and Gag-specific memory precursor CD8⁺ T cells exhibited distinct cellular states. Here, we defined memory precursor cells as vaccine-induced cells that are detected at M7 that remained detectable at the M24 durability time-point. Env- and Gag-specific CD8⁺ T cells were identified within the REFLEX dataset by matching participant-specific TCR sequences to the experimentally defined TCRα or TCRβ libraries from Figure 1. Projection of these cells onto the CD8⁺ T cell UMAP revealed substantial phenotypic overlap between Env- and Gag-specific responses (Fig. 5, A and B). However, Gag-specific CD8⁺ T cells were enriched within the cycling/proliferative population compared with Env-specific CD8⁺ T cells (Fig. 5C).

**Fig. 5.**
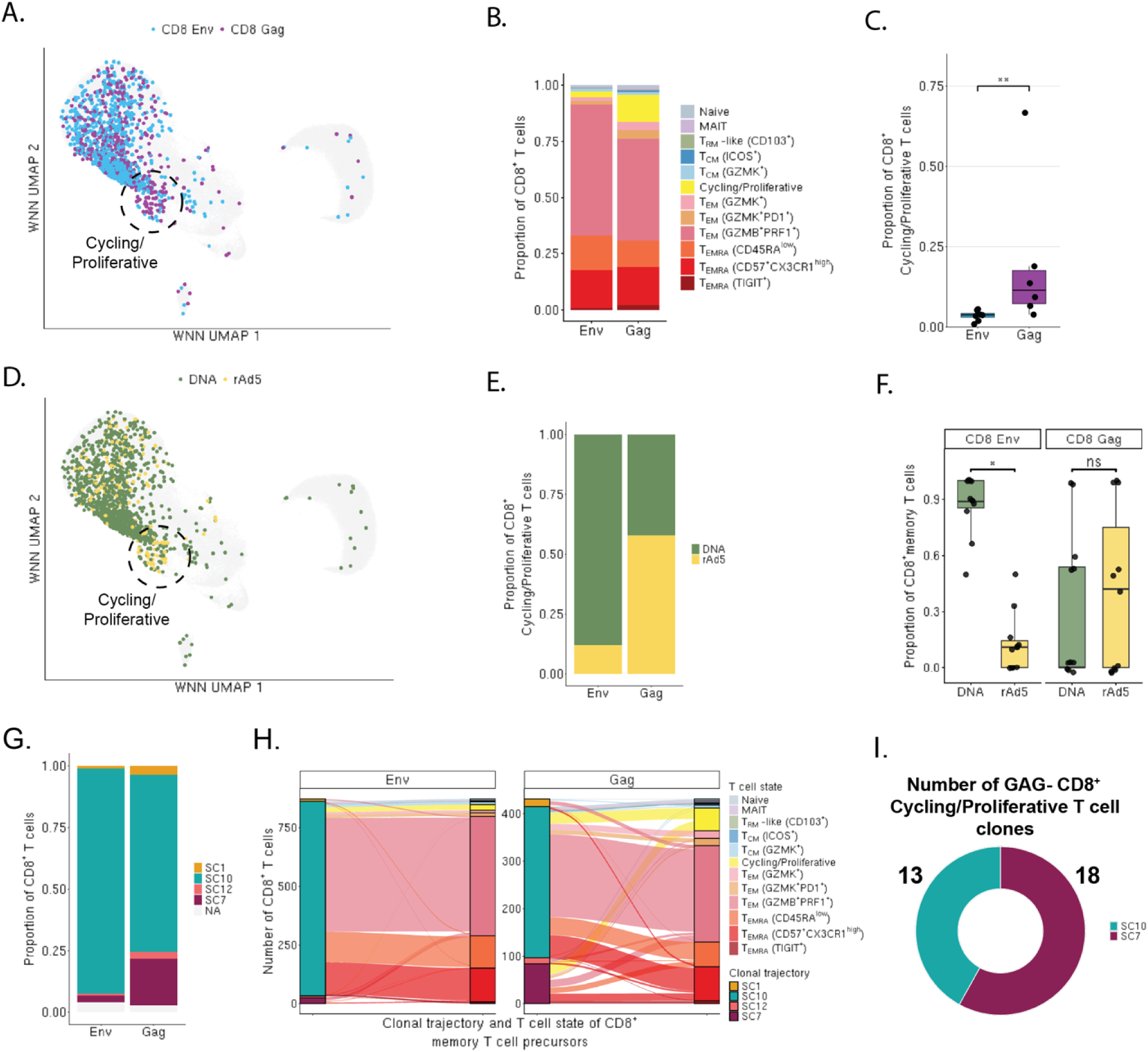
Distinct clonal trajectories and cellular states of Env- and Gag-specific CD8⁺ memory precursor T cells **A)** UMAP projection of antigen-specific CD8⁺ T cells identified by matching participant-specific TCR sequences to experimentally defined Env- and Gag-specific TCRα/TCRβ libraries, colored by antigen specificity **B)** Stacked bar plot showing the distribution of annotated CD8⁺ T cell states among Env- and Gag-specific CD8⁺ T cells **C)** Participant-level quantification of the proportion of Env- and Gag-specific CD8⁺ T cells in the cycling/proliferative state. Each point represents an individual participant. Statistical comparisons were performed using paired Wilcoxon signed-rank tests **D)** UMAP projection of Env- and Gag-specific CD8⁺ T cells colored according to the vaccine dose at which the corresponding clone was first detected, following DNA prime or rAd5 boost **E)** Proportion of Env- and Gag-specific cycling/proliferative CD8⁺ T cells derived from clones first detected following DNA prime or rAd5 boost **F)** Proportion of Env- and Gag-specific memory clones induced by DNA prime or rAd5 boost. Memory clones were defined as vaccine-induced clones detected at M7 that remained detectable at M24. Each point represents an individual participant and antigen-specific response. Statistical comparisons were performed using paired Wilcoxon signed-rank tests with BH correction for multiple comparisons **G)** Distribution of longitudinal clonal trajectory clusters among Env- and Gag-specific memory clones detected at M24. Memory clones were defined as vaccine-induced clones detected at M7 that remained detectable at M24 **H)** Alluvial plot showing the relationship between longitudinal clonal trajectory and CD8⁺ T cell state among memory precursor cells. Flows represent the number of cells connecting each trajectory cluster with its corresponding T cell state. Data are shown separately for Env- and Gag-specific responses **I)** Proportion of unique clones within the Gag-specific cycling/proliferative population assigned to each longitudinal clonal trajectory cluster. Clones were defined using paired TCRα and TCRβ sequences when both chains were available and by the available chain when only one chain was detected *\* P* < 0.05; ** *P* < 0.01; ns, not significant

This cycling/proliferative phenotype was predominantly associated with rAd5-induced Gag-specific CD8⁺ T cells (Fig. 5, D and E, fig. S8B). Notably, rAd5-induced cells comprised a greater proportion of the Gag-specific memory compartment, whereas Env-specific memory remained predominantly composed of DNA prime-induced cells (Fig. 5F). Among M7 vaccine-induced clones that persisted to M24, Env-specific memory precursor clones were predominantly associated with trajectory cluster SC10, whereas a subset of Gag-specific memory precursor clones localized to SC7, a trajectory enriched for cycling/proliferative T cells (Fig. 5G).

Linking clonotypes to their cellular states further revealed that rAd5-induced Gag-specific cells exhibiting SC7 kinetics contributed to the cycling/proliferative memory precursor population that persisted through M24 (Fig. 5H). This population was not observed among Env-specific memory precursors. Consistent with the greater contribution of rAd5-induced clones to Gag-specific memory, SC7-derived clones comprised a larger fraction of Gag-specific cycling/proliferative population than DNA prime-induced clones (Fig. 5I).

Collectively, these findings link the greater representation of cycling/proliferative Gag-specific CD8⁺ T cells to rAd5-induced clonal trajectories that persist into long-term memory. Thus, the DNA/rAd5 vaccine regimen differentially shaped the clonal origins and cellular states of Env- and Gag-specific memory precursors CD8⁺ T cells.

## DISCUSSION

In this study, we combined antigen-specific TCR identification with longitudinal TCR sequencing and single-cell multi-omics to characterize HIV-1 vaccine-induced T cell responses following DNA/rAd5 prime-boost vaccination. We found that Env- and Gag-specific CD8⁺ T cell responses followed distinct clonal trajectories after DNA prime and rAd5 boost, resulting in differences in repertoire composition, cellular state, and long-term persistence. Env-specific responses were characterized by greater contribution of DNA prime-induced clones to durable memory, whereas Gag-specific responses included a greater contribution of rAd5-induced clones and a persistent cycling/proliferative CD8⁺ T cell population.

We applied a two-step enrichment and expansion strategy to generate high-confidence HIV-1 Env- and Gag-specific CD4⁺ and CD8⁺ TCR libraries and demonstrated that TCR-derived response magnitudes were generally concordant with those measured by ICS. The greatest discordance occurred for Gag-specific CD8⁺ T cell responses, which were detected at higher frequencies by TCR sequencing. This difference likely reflects the distinct features measured by the two approaches: ICS detects cells that produce detectable cytokines following *in vitro* antigen stimulation, whereas TCR sequencing captures antigen-specific clonotypes independent of the capacity of individual cells produce cytokines under these conditions (*51*). The enrichment of Gag-specific CD8^+^ T cells within the cycling/proliferative state observed by REFLEX may provide one potential explanation for this discordance. Thus, TCR sequencing provides a complementary measure of vaccine-induced cellular immunity that can capture antigen-specific populations alongside bulk cytokine-based assays.

Beyond quantifying vaccine responses, the generation of high-confidence Env- and Gag-specific TCR libraries provides a scalable resource for tracking vaccine-induced T cell responses across time and cohorts. As additional HIV-1 vaccine candidates progress through pre-clinical and clinical development, expanding such antigen-specific TCR catalogues could facilitate longitudinal monitoring of vaccine responses and identification of clonal features associated with protective immunity.

The DNA prime and rAd5 boost differentially shaped the clonal composition and persistence of Env- and Gag-specific CD8⁺ T cell responses. More Env-specific than Gag-specific CD8⁺ T cell clones were detected after DNA prime, establishing a broader Env-specific repertoire before rAd5 boost. Although rAd5 recruited newly detected Env- and Gag-specific clones at similar frequencies, the Env-specific repertoire subsequently became more clonal, consistent with preferential expansion and persistence of previously primed clones. In contrast, rAd5-induced clones contributed more substantially to the Gag-specific memory compartment. This suggests that the broader Env-specific repertoire established during prime provides a larger pool of clones available for recall and long-term persistence. Multi-clade and mosaic DNA vaccines can generate broader and more cross-reactive T cell responses, raising the possibility that the greater diversity of Env-specific clones established during prime contributed to their preferential persistence following rAd5 boost (*52–54*). However, these difference may reflect intrinsic antigen properties and differences in vaccine design and dosing, including the multi-clade Env and monovalent Gag immunogens and the 3:1 ratio of rAd5 vectors encoding Gag relative to Env (*21*).

Integration of longitudinal TCR tracking with single-cell REFLEX analyses further connected clonal history with the cellular states of Env- and Gag-specific memory precursors. Env-specific memory precursors were predominantly derived from DNA prime-induced clones that subsequently expanded following the rAd5 boost and were enriched within the SC10 clonal trajectory and differentiated cytotoxic *GZMB^+^PRF1^+^* T_EM_ state. In contrast, rAd5-induced Gag-specific clones were more frequently associated with the SC7 trajectory and a highly activated cycling/proliferative state that remained detectable at M24 and contributed to the durable Gag-specific memory compartment. Altogether, differences in the timing of clonal recruitment were associated with distinct cellular states among Env- and Gag-specific memory precursors, although these differences may partly reflect the interval between clonal induction and sampling. Nevertheless, the cellular state in which these clones persist may influence their functional capacity upon subsequent antigen encounter.

Our study has several limitations. First, single-cell multi-omics profiling was performed at a singular time-point (M7), limiting our ability to determine how memory precursor states evolve into long-term memory or to distinguish effects of vaccine platform and antigen from differences in the timing of clonal induction. Although longitudinal TCR tracking identified clones that persisted to M24, we cannot ascertain whether their M7 transcriptional and phenotypic states were maintained or remodeled over time. Second, the limited number of participants and antigen-specific cells restricted our ability to quantitatively assess differences between DNA- and rAd5- induced Env- and Gag- specific CD8⁺ T cells or their functional consequences. Finally, because all participants remained HIV-1-negative throughout follow-up, we could not directly associate the identified clonal, phenotypic or molecular features with protection from HIV-1 acquisition. Future studies incorporating longitudinal single-cell profiling and functional assessment of vaccine-induced T cells, particularly in cohorts with and without HIV-1 acquisition, will be needed to determine whether these features can represent correlates of protective immunity Our study provides several methodological advances for the characterization of antigen-specific T cell responses. First, we developed an enrichment and expansion strategy capable of generating high-confidence antigen-specific TCR libraries without prior knowledge of relevant epitopes. Second, we applied an unbiased longitudinal trajectory framework to characterize clonal expansion and persistence overtime. Third, we used REFLEX, a single-cell multi-omics platform that simultaneously captures TCR sequences, transcriptomic profiles, and surface protein expression. Together, these approaches enabled an integrated characterization of HVTN 505 vaccine-induced Env- and Gag-specific T cell responses across clonal dynamics, cellular phenotype, and molecular state, providing a comprehensive framework for evaluating vaccine-induced cellular immunity.

## MATERIALS AND METHODS

### Study design

*Clinical Cohort:* HVTN 505 (ClinicalTrials.gov Identifier: NCT00865566) was a multicenter, double-blind, randomized, placebo-controlled phase 2b trial that evaluated the efficacy of a multiclade vaccine research center (VRC) DNA/rAd5 HIV-1 vaccine regimen for HIV-1 prevention (*21*). The study enrolled 2,504 HIV-1–, HIV-2–, and Ad5-seronegative men and transgender women aged 18 to 50 years in the United States. All participants provided informed consent. Participants were randomized to receive three doses of DNA vaccine, administered 4 weeks apart, consisting of Gag, Pol, and Nef from HIV-1 clade B and Env from clades A, B, and C, or phosphate-buffered saline (PBS) placebo. Sixteen weeks after the final DNA dose, participants received a recombinant adenovirus serotype 5 (rAd5) boost consisting of four vectors expressing Gag-Pol from HIV-1 clade B and Env from clades A, B, and C, or placebo, administered at a ratio of 3:1:1:1. Participants were scheduled for follow-up visits at months 0, 1, 2, 2.5, 6, 7, and 9 and every 3 months thereafter through 24 months. Cryopreserved PBMC samples were available at baseline (M0), M2.5, M7, M12, and M24.

*Study participants:* A sub-cohort of randomly selected participants who received all scheduled vaccine injections and remained HIV-1 negative through M24 was previously identified by the HVTN for follow-up immunogenicity analyses. From this sub-cohort, 16 vaccinated participants were selected for the present study based on sufficient sample availability at all longitudinal time points and detection of at least one Env- or Gag-specific T cell response by ICS.

### TCR library generation and sequencing

RNA was extracted using the NucleoSpin RNA Plus kit (Macherey-Nagel) from all sorted cytokine-positive and cytokine-negative cells, as well as from 0.5x10^6^ Env- and Gag-expanded T cells and 5x10^6^ total PBMCs collected at M0, M2.5, M7, M12, and M24 for all participants. TCRα/β sequencing libraries were prepared using the SMARTer Human TCRα/β Profiling Kit (Takara) according to the manufacturer’s protocol. Final TCR libraries were quantified using a Qubit Fluorometer (Invitrogen) and assessed using a TapeStation (Agilent). Cytokine^+^ and cytokine^-^ TCR libraries were pooled, whereas expanded T cell TCR libraries were pooled with longitudinal PBMC TCR libraries, and the two pools were sequenced separately using an Illumina NextSeq 1000/2000 P3 300-cycle sequencing flow cell and cartridge (Illumina). Library pools were sequenced at 650 pM with 10% PhiX (Illumina). Sequencing data were processed using MiXCR v4.3.2 (*84*).

### TCR trajectory analyses and identification of vaccine-specific trajectory clusters

The 1,500 most abundant TCR clones at each time point for each participant were selected, and longitudinal clone frequencies were normalized to the peak frequency of each clone as previously described (*6*). Non-detected clones were assigned a frequency equal to one-half the minimum detected frequency at the corresponding time point for each participant. Hierarchical clustering was performed using Ward’s method on Euclidean distances calculated from log10-transformed normalized longitudinal frequencies. Multiple cluster resolutions were evaluated, and a 10-cluster resolution was selected for downstream analyses. UMAP was performed using the first five principal components calculated from the normalized longitudinal frequencies. Enrichment of antigen-specific TCRs within trajectory clusters was assessed using Fisher’s exact tests, with BH-adjusted *P* < 0.05 considered significant. Env- and Gag-specific TCRs were identified in longitudinal datasets by exact matching of TCR sequences within participants. Longitudinal trajectory analyses were performed separately for TCRβ and TCRα chains.

### Statistical analyses

Statistical analyses were performed in R v4.4.0. For comparisons between two groups, Wilcoxon signed-rank tests were used for paired samples and Wilcoxon rank-sum tests for unpaired samples, as indicated in the figure legends. Spearman rank correlation was used to assess concordance between TCR sequencing– and ICS-derived response magnitudes. Vaccine-enriched TCR clones were identified using Fisher’s exact tests, with *P* values adjusted for multiple comparisons using the BH procedure. For analyses involving multiple pairwise comparisons, *P* values were similarly adjusted using the BH procedure to control the false discovery rate. Unless otherwise indicated, statistical tests were two-sided, and *P* < 0.05 was considered statistically significant for individual comparisons; BH-adjusted *P* < 0.05 was considered statistically significant for analyses involving multiple comparisons.

## List of Supplementary Materials

Materials and Methods Fig. S1 to Fig. S8 Table S1 to Table S8 References (85–86)

## Acknowledgments

We gratefully acknowledge all participants who enrolled in the HVTN 505 clinical trial and all members and staff who contributed to its conduct. This study was made possible by samples provided by the HVTN as part of HVTN auxiliary study number 324_EXS_Newell_505. We thank Holly Jane for her advice regarding this study, David Koelle for his experimental guidance, the Vaccine Research Center (VRC) at the National Institute of Allergy and Infectious Diseases (NIAID) for providing the HIV-1 Env and Gag Clade B peptide pools, and the Fred Hutch Flow Cytometry Core for their technical support. Figures were created using Illustrator (Adobe) and BioRender.

## Author contributions

Conceptualization: GX, KMB, DRG, AAM, STK, MES, PGT, SCD, AFG, EWN Methodology: GX, KMB, AAM, MRH, PGT, PJS, SCD, AFG, EWN

Investigation: GX, JXY, SMI, SNK, EM Visualization: GX

Formal analysis: GX

Funding acquisition: PJS, SCD, MJM, AFG, EWN Supervision: EWN

Writing – original draft: GX

Writing – review & editing: GX, SNK, KMB, AAM, DRG, MRH, SCD, AFG, EWN

## Data and materials availability

Detailed experimental protocols are available upon request. All data and code are available at: https://doi.org/10.5281/zenodo.22308605

