## Supplementary Materials for "HIV-1 prime-boost vaccination shapes distinct clonal trajectories and memory precursor states of Env- and Gag-specific T cells"

**Supplementary materials and methods**

**Human T cell isolation and Expansion***:*

*Peptide stimulation and magnetic enrichment using AIM markers:*

For generation of HIV-1 Env- and Gag-specific TCR libraries, M7 PBMCs were thawed and rested overnight at 2x10^6^ cells/mL in R10 medium [RPMI-1640 (Gibco) supplemented with 10% heat-inactivated fetal bovine serum, 1% penicillin-streptomycin (Gibco), and 0.5% L-glutamine (Gibco)] in T25 culture flasks. Cells were then stimulated for 18 hours at 5x10^6^ cells/mL in 96-well plates with vaccine-matched HIV-1 Env or Gag peptide pools (1μg/mL) in the presence of anti-CD40 blocking antibody (1μL/mL; Miltenyi Biotec, 130-094-133) and anti-CD154-PE antibody (1:200; clone TRAP1, BD Biosciences). Peptide pools were provided by the Vaccine Research Center (VRC) and consisted of 15-mer peptides overlapping by 11 amino acids.

Following stimulation, cells were washed with MACS buffer and stained with anti-CD137-PE (2μL per 1x10^7^ cells; clone 4-1BB, Miltenyi Biotec) and anti-OX40-PE (2μL per 1x10^7^ cells; clone ACT35, Miltenyi Biotec) in a total volume of 50μL for 10 min at 4°C. Cells were subsequently washed and incubated with anti-PE MicroBeads (20μL per 1x10^7^ cells; Miltenyi Biotec) in a total volume of 100μL for 15 min at 4°C. CD137⁺, CD154⁺, and/or CD134⁺ activation-induced marker (AIM) cells were enriched by positive selection using LS columns (Miltenyi Biotec) according to the manufacturer's instructions.

*Primary T cell expansion and culture*

Following AIM enrichment, cells were rested for at least 2 hours at 37°C in CTL medium in 24-well plates before initiation of the canonical ex vivo T cell rapid expansion protocol (REP) (*38*). Enriched cells were transferred to T25 culture flasks containing 5mL of REP master mix. REP master mix consisted of 30x10^6^ irradiated PBMCs, 5x10^6^ irradiated TM-LCLs, 30 ng/mL anti-CD3 antibody (OKT3), and 1μg/mL anti-CD28 antibody (BioLegend) in CTL medium, prepared at a final volume of 25mL. IL-2 and IL-15 were added to a final concentration of 50U/mL each beginning 1 day after initiation of expansion. Cell counts were performed daily beginning on day 5. Half-medium changes with fresh IL-2 and IL-15 were performed every other day when cultures did not require splitting. Cultures were split when cell concentrations exceeded 1.5x10^6^ cells/mL. Expanded cells were cryopreserved in freeze-down medium on day 15.

**Generation of HIV-1 Env and Gag TCR library**

*Peptide re-stimulation & cytokine capture*

Autologous antigen-presenting cells (APCs) were generated for each participant by thawing PBMCs, depleting T cells using the Pan T Cell Isolation Kit (Miltenyi Biotec), and labeling the remaining cells with CellTrace Violet (Thermo Fisher Scientific). Cells were rested overnight at 2x10^6^ cells/mL in R10 medium supplemented with 500U/mL IL-4 and 500U/mL GM-CSF. On the same day, Env- and Gag-expanded T cells were thawed and rested overnight at 2x10^6^ cells/mL at 37°C in CTL medium. Following overnight rest, autologous APCs were pulsed for 30mins with vaccine-matched Env or Gag overlapping peptide pools (1μg/mL) or DMSO control in the presence of anti-CD40 blocking antibody (1μL/mL; Miltenyi Biotec, 130-094-133). Env- or Gag-expanded T cells were then added to the corresponding APCs at a 2:1 T cell-to-APC ratio and stimulated for 4 hours at 37°C.

Following stimulation, IFNγ, TNFα, and IL2 secretion was assessed using the cytokine secretion assay (Miltenyi Biotec). IFN-γ-FITC, TNF-α-APC, and IL-2-PE cytokine secretion reagents were combined at an 1:1:1 ratio according to the manufacturer's instructions. Cells were subsequently stained with anti-CD3-BUV395 (5:200; clone UCHT1, BD Biosciences), anti-CD4-BUV805 (5:200; clone SK3, BD Biosciences), anti-CD154-BV711 (2:200; BioLegend), anti-CD8-APC-H7 (2:500; clone SK1, BD Biosciences), and 7-AAD (BD Pharmingen). CD4⁺ and CD8⁺ T cells were sorted separately by flow cytometry into cytokine^+^ and cytokine^-^ populations. Cytokine^+^ cells were defined based on expression of IFNγ^high^/IL2^+^ or TNFα^+^/CD154^+^ expression, whereas cytokine^-^ cells lacked detectable IFNγ, IL2, TNFα, and CD154 secretion (See Figure S1A). Sorted populations were subsequently used for TCR sequencing.

*Identification of candidate antigen-specific T cell clones*

Following TCR sequencing, candidate HIV-1 Env- and Gag-specific TCR clones were identified by comparing clonal frequencies across the cytokine^+^, cytokine^-^, expanded T cells, and M7 PBMC populations. A clone was considered a candidate antigen-specific TCR if it met all of the following criteria: (i) its frequency was at least 4-fold higher in the expanded T cell population than in M7 PBMCs; (ii) its frequency was at least 2.5-fold higher in the cytokine^+^ population than in the expanded T cell population; (iii) its frequency was higher in the cytokine^+^ population than in the corresponding cytokine^-^ population; and (iv) the clone had an unique molecular identifier (UMI) **count <2 at M0**. Candidate antigen-specific TCR clones identified using these criteria were subsequently matched by exact TCR sequence within participant to the longitudinal bulk TCR dataset to identify HIV-1 Env- and Gag-specific CD4⁺ and CD8⁺ T cell clones.

**TCR repertoire analyses**

Clonal breadth was defined as the number of unique antigen-specific clones detected per participant, where clones were defined by the CDR3 amino acid sequence together with the corresponding V- and J-gene usage. For each clone, its clonal frequency was calculated separately for every longitudinal PBMC sample in which the clone was assessed. Clonal frequencies were calculated as the sum of UMI counts for the clone divided by the total UMI counts for the sample. For each participant, cell type, antigen, and time point, TCR response magnitude was calculated as the sum of the frequencies of all experimentally identified antigen-specific clones that passed the filtering criteria states above. TCR repertoire diversity was quantified using Shannon entropy, calculated from the relative frequency distribution of clones within each sample. Clonal richness was defined as the number of unique clones detected within each sample. Clonal evenness was measured using Pielou’s evenness index, calculated as Shannon entropy divided by the natural logarithm of clonal richness. Clonality was calculated as 1 - Pielou’s evenness.

**Identifying DNA- and rAd5- induced clones**

Clones were classified according to their first detectable expansion following vaccination. DNA prime-induced clones were defined as clones that were detectable at M2.5, following the DNA prime, and had a higher clonal frequency at M2.5 than at baseline (M0). rAd5 boost-induced clones were defined as clones that were not detectable at M2.5 and subsequently had a higher clonal frequency at M7, following the rAd5 boost, than at both M0 and M2.5. Clones that did not meet these criteria were not assigned to either group.

**Calculation of clonal decay**

Clones detected at M7 were tracked longitudinally per participant. Clones not detected at M12 were assigned a frequency equal to half of the smallest non-zero clonal frequency observed in the corresponding M12 sample to allow for logarithmic transformation. Clonal decay was quantified as $\left( \ln\left( f_{M12} \right)-\ln\left( f_{M7} \right) \right)/\left( M12-7 \right)$, where $f_{M7}$and $f_{M12}$denote clone frequencies at M7 and M12, respectively, with more negative slopes indicating faster contraction of clonal abundance over time.

**Identification of antigen-specific TCRs through statistical enrichment tests**

Fisher’s exact tests were used to identify clones that were significantly enriched following vaccination. Clonal counts at M2.5 and M7 were independently compared with baseline (M0) to identify clones significantly enriched following the DNA prime or rAd5 boost, respectively. Clonal counts at M7 were also compared with M2.5 to identify clones significantly enriched following the rAd5 boost relative to the post-prime repertoire. For each comparison, the Fisher’s exact test evaluated the enrichment of individual clones based on their UMI counts relative to the remaining sequenced repertoire. Clones were considered significantly enriched if they met both of the following criteria: (1) a Benjamini–Hochberg false discovery rate (FDR)-adjusted P value < 0.05 and (2) log2 fold-change > 2 in clonal abundance between the comparison time points. Clones meeting these criteria were classified as vaccine expanded.

**Longitudinal intracellular cytokine staining (ICS)**

ICS was performed on the same vial of PBMC sample used above for longitudinal TCR sequencing. Cells used for ICS were rested overnight in R10 media at 37°C after PBMC were thawed. Cells were then stimulated with vaccine-matched overlapping peptide pools for Env- and Gag- or DMSO for 6h in the presence of 1X Brefeldin A (Biolegend) and 1X Monensin (Biolegend) at 37°C. Cells were then stained with Live/Dead fixable blue cocktail (Invitrogen) with Human TruStain FcX (Biolegend) for 20mins at room temperature (RT) prior to a 10mins fixation with 1X FACS Lyse (BD Bioscience) and 20mins permeabilization with 1X FACS Perm (BD Bioscience) at RT. Cells were then stained with antibody cocktail consisting of anti-CD3-BUV395 (0.4/50, clone UCHT1, BD Biosciences), anti-CD8-BUV805 (0.05:50, clone SK1, BD Biosciences), anti-CD4-BV480 (0.1:50, clone SK3, BD Horizon), anti-IFNγ-V450 (0.25:50, clone B27, BD Horizon), anti-TNF-α- FITC (0.13:50, clone MAb11, eBioscience), anti-IL2 -PE (1.5:50, clone MQ1-17H12, BD Pharmingen) and anti-CD154 -APC (4:50, clone TRAP1, BD Pharmingen), for 30mins at RT. All cells were acquired on BD FACSymphony A3 cell analyzer (BD Biosciences). Data were analyzed using FlowJo v10.8 software (BD Biosciences) and all results were DMSO background subtracted.

**Single cell sequencing using REFLEX:**

*Sample preparation for REFLEX*

11 participants were selected for single-cell sequencing based on their Env- and Gag- specific T cell responses as determined by TCR sequencing. We selected for participants with detectable responses to both antigens. PBMC samples at M7 for the selected participants were thawed and pan-T cell isolation (Miltenyi Biotec) was performed to isolate for CD3^+^ T cells. Cells were then stained with Human TruStain FcX (Biolegend) for 10mins at 4°C, washed and then stained with a custom TotalSeq-C antibody cocktail (table S7) for 30mins at 4°C. The antibody cocktail was filtered through a 0.1uM Ultrafree-centrifugal filter (Millipore Sigma) prior to stain. Cells were washed twice with Dulbecco’s Phosphate Buffered Saline (DPBS) (Gibco) prior to fixation.

*Single cell sequencing of ADT, GEX, and TCR libraries with REFLEX*

REFLEX was performed using the GEM-X v1 version of the protocol with the inclusion of resolvable oligos for the detection of TotalSeq C antibodies as previously described^51^. Additional protocol details can be found at [protocols.io](https://www.protocols.io/view/reflex-on-gem-x-v1-e6nvww797vmk/v1). scRNAseq FLEX libraries were sequenced using a NextSeq2000 P4 300 cycle flow cell, with the following parameters: Read1:80 cycles, Read2:200 cycles. A read depth of 5000 reads per cell for TCR and ADT libraries or 10,000 reads per cell for GEX libraries was targeted. REFLEX TCR alignment was performed using MIXCR’s (v4.7) “generic-ht-single-cell-amplicon workflow”. Sample barcodes were provided using the –-sample-sheet parameter to demultiplex the TCR libraries. Cell barcode alignment used the 10X barcode whitelist in the –set-whitelist parameter. 10X 5’ VDJ fastq alignment was performed using MIXCR’s (v4.7) “10x-sc-xcr-vdj” preset.  The resulting clones.tsv files from both workflows were integrated into their respective Seurat objects using scRepertoire (v2.5.3).

*Single-cell sequencing analyses*

Quality control, data integration and analyses were performed using Seurat (*85*). Seurat SCTransform was used to integrate our two sequencing runs together. CD4 and CD8 ADT expression were compared between cells and double positive CD4 and CD8 cells were removed prior to weighted-nearest neighbor (WNN) clustering (*86*). Non-T cell clusters were removed based on their low RNA expression of *CD3D* and *CD3E,* and low ADT expression of CD3, and high ADT expression of CD10 and CD16. CD8 and CD4 T cells were then re-clustered separately using WNN analyses after separating the two-cell types based on their ADT expression. Elbow plots were generated to determine the optimal number of PCs to use for RNA and ADT multi-modal WNN clustering. Cells were clustered using different clustering resolutions and cell type annotations were determined based on RNA and ADT expression. ADT positivity thresholds for each marker were determined using a two-component Gaussian mixture model fitted to the normalized ADT expression distribution across all cells, with the threshold defined as the midpoint between the inferred negative and positive populations. From this 8 CD4⁺ T cell clusters and 12 CD8⁺ T cell clusters were defined.

*Single cell longitudinal clustering*

The top 1500 most abundant TCRβ and TCRα clones from above were filtered to only include clones that are detected within our single-cell dataset. The longitudinal TCRβ and TCRα datasets were filtered separately prior to being combined for unbiased trajectory clustering analyses. For clones that were found in both datasets, the TRB clonal frequencies were kept. Clones were matched within participant by CDR3 amino-acid sequence and V and J-gene. Single cell longitudinal trajectories were re-clustered and determined as described above.

*Differential expression analyses*

DEGs and DEPs were identified for each single-cell trajectory clonal cluster using a one-versus-all approach, where each cluster was compared against all remaining cells using Wilcoxon rank-sum testing. For RNA analysis, genes with an adjusted *P*-value <0.05 and average log2 fold-change >0.5 were considered significantly enriched, and the top enriched genes from each trajectory cluster were selected for visualization. For ADT analysis, differentially abundant surface proteins were identified using the same statistical framework, with markers considered significant at an adjusted *P*-value <0.05; the most enriched proteins were selected based on ranked average log2 fold-change values.

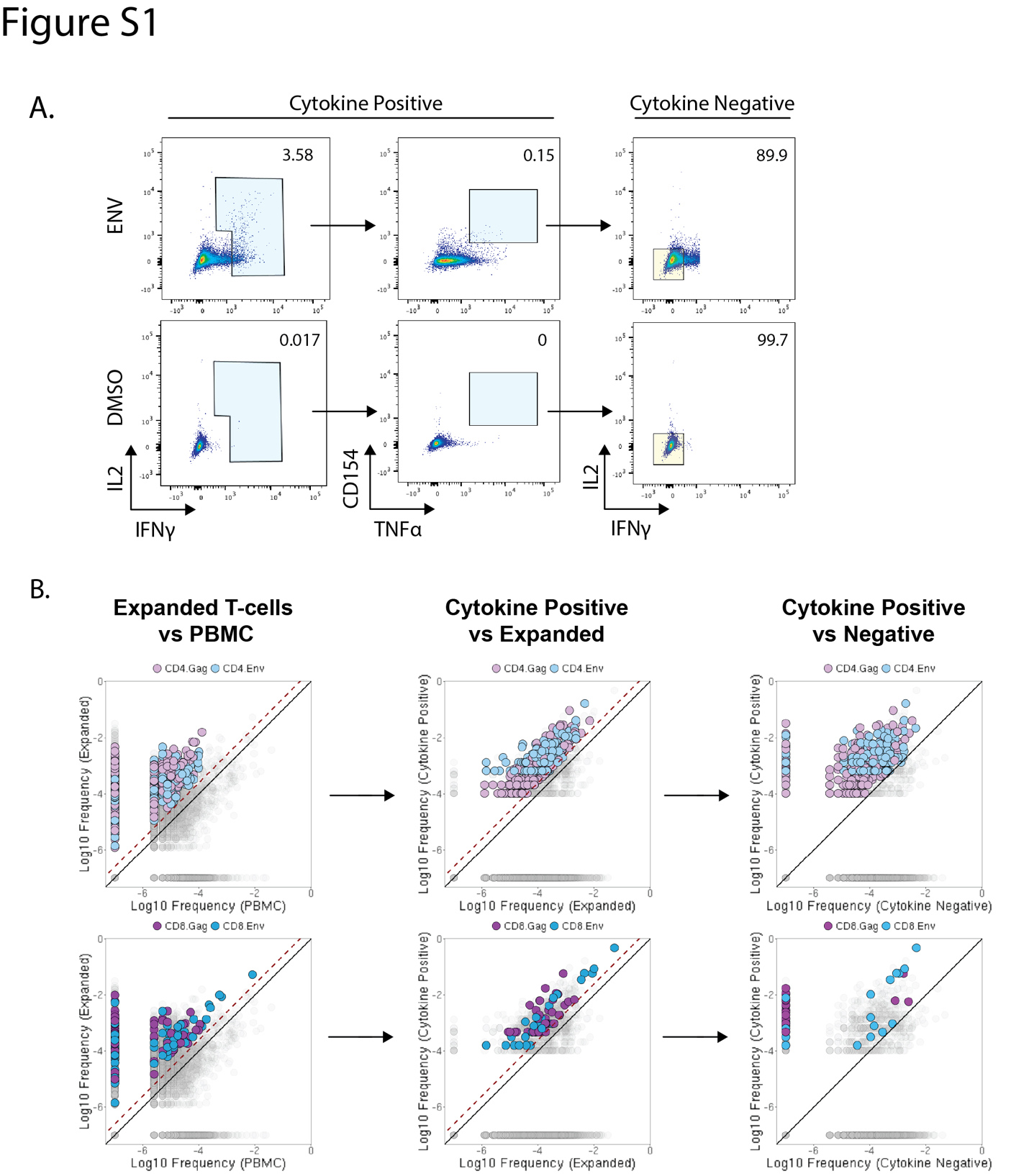

**Fig. S1. Identification of antigen-specific TCRs following cytokine-based enrichment and bulk TCR sequencing**

A) Gating strategy used to isolate Env- and Gag-specific cytokine-positive and cytokine-negative CD4⁺ and CD8⁺ T cells following peptide restimulation. The blue box indicates the pooled population used for sorting cytokine^+^ cells, and the yellow box indicates the population used for sorting cytokine^-^ cells
B) Filtering strategy used to identify antigen-specific TCRs following cytokine^+^ and cytokine^-^ cell sorting and bulk TCR sequencing. Each dot represents a unique TCRβ clone, with clonal frequencies compared between the indicated conditions. Sequential frequency-based filters were applied to identify clones enriched following T cell expansion, further enriched in the cytokine^+^ population, and preferentially represented in the cytokine^+^ versus cytokine^-^ population. Clones meeting all filtering criteria were classified as antigen specific. Colored dots indicate cytokine^+^ clones that met all filtering criteria, whereas gray dots represent all other TCRβ clones detected in the sequenced repertoire. See Supplemental Materials and Methods for detailed filtering criteria

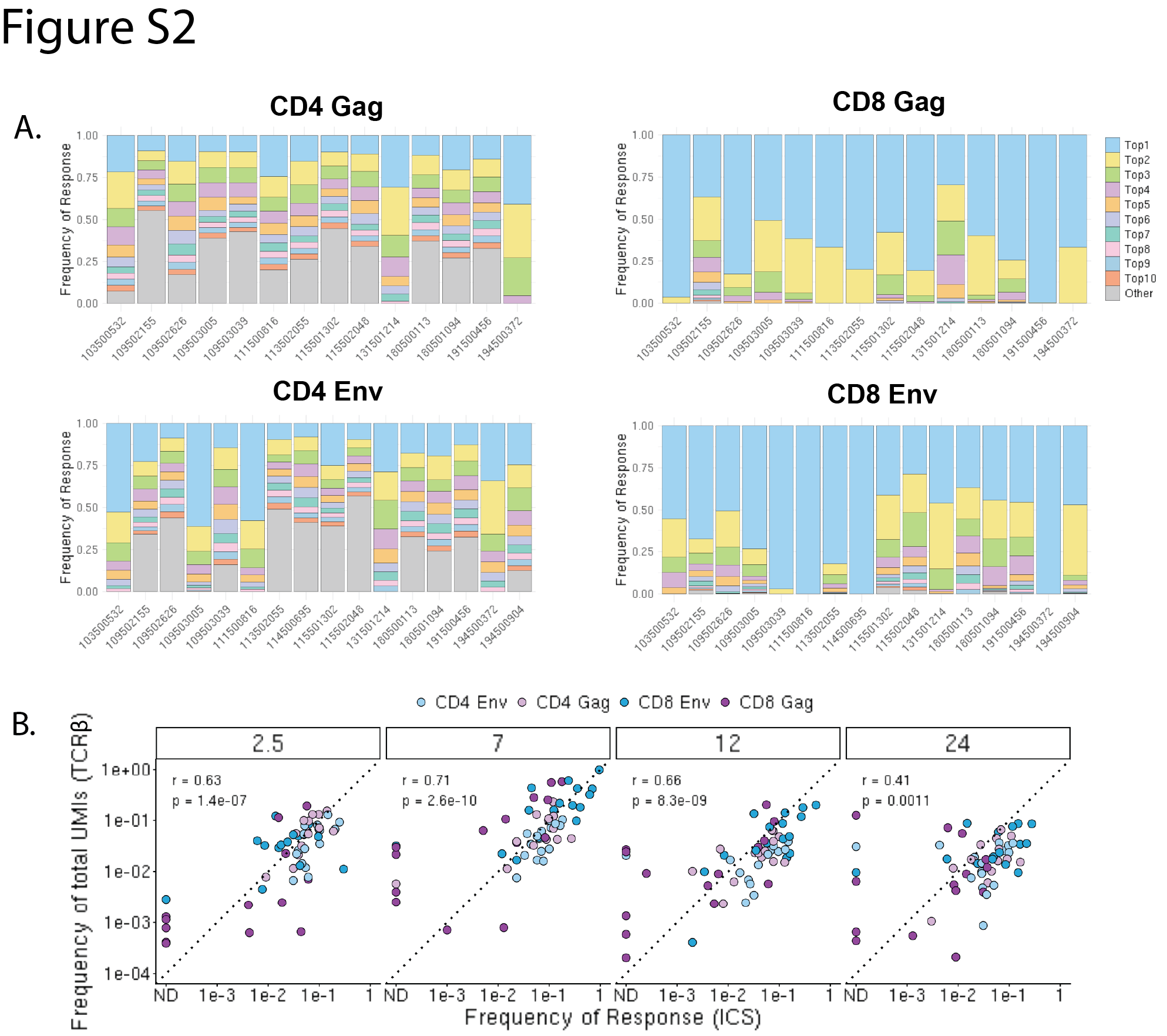

**Fig. S2. Comparison of vaccine-specific TCR and cytokine responses**

A) Stacked bar plots showing the top 10 TCRβ clones for each participant, cell type, and antigen
B) Scatter plots showing the correlation between vaccine-specific TCRβ and ICS response magnitudes across participants for Env- and Gag-specific CD4⁺ and CD8⁺ T cell responses at each study visit. Each point represents an individual participant. Spearman rank correlations were calculated separately for each time point and cell type–antigen combination. Response magnitudes are shown on logarithmic scales, with the Spearman correlation coefficient and P-value indicated

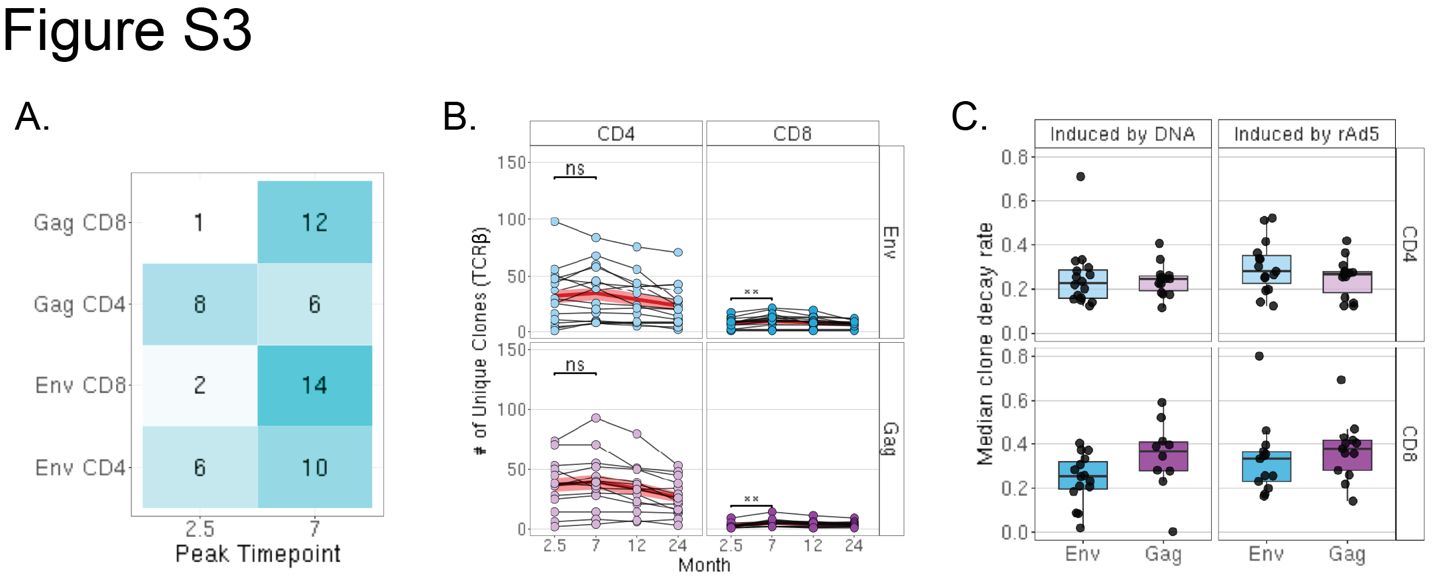

**Fig. S3. Longitudinal dynamics of vaccine-induced T cell repertoire**

A) Heatmap showing the distribution of peak response time points across antigen-specific T cell subsets. The number of participants reaching their peak response at each time point is indicated within each cell. Differences in peak time point distributions were assessed using Fisher’s exact test with *P*-values adjusted using the BH method (P = 0.014)
B) Longitudinal changes in TCRβ repertoire breadth assessed by clonal richness, defined as the number of unique clones detected within each antigen- and T cell subset per participant. Red lines indicate the mean clonal richness across participants at each time point, and shaded ribbons indicate the standard error of the mean (SEM). Differences between groups were assessed using Wilcoxon rank-sum tests with *P*-values adjusted using the BH method
C) Boxplots showing participant-level median clonal decay rates for the indicated antigen and T cell subsets. Clonal decay rates were calculated from longitudinal changes in clone frequency between M7 and M12 among clones with detectable frequencies at M7

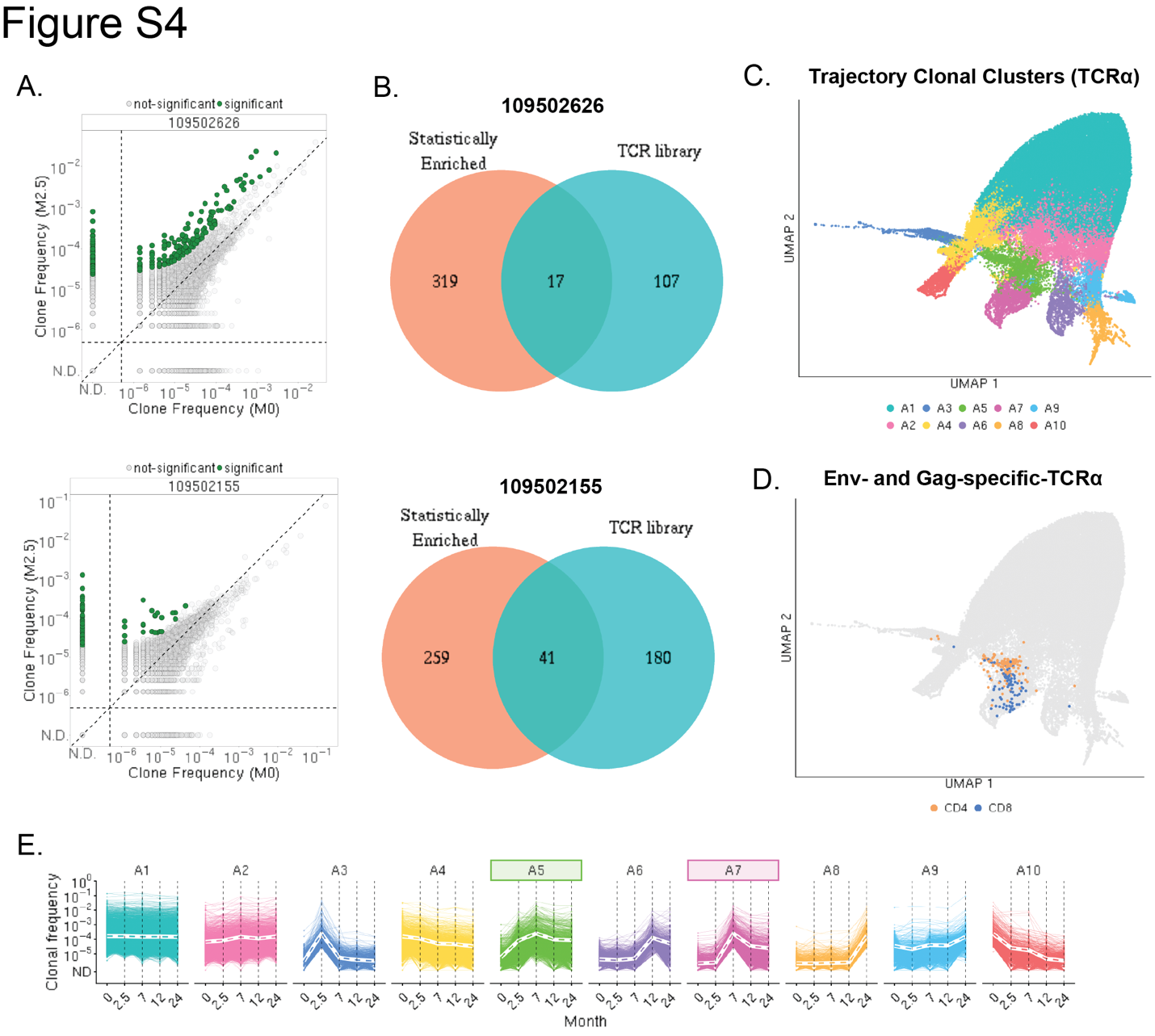

**Fig. S4. Longitudinal clonal trajectories analyses**

A) Biaxial plots comparing TCRβ clonal frequencies at M0 and M2.5 for two representative participants. Clones identified as statistically enriched at M2.5 based on a >2-fold increase and FDR-adjusted P < 0.05 by Fisher’s exact test are shown in green
B) Venn diagram showing the overlap between clones identified by statistical enrichment at M2.5 and experimentally identified antigen-specific TCRβ clones
C) UMAP representation of longitudinal TCRα clonal trajectories for the top clones, colored according to trajectory clusters identified by hierarchical clustering
D) UMAP colored by experimentally identified CD4⁺ (orange) and CD8⁺ (blue) vaccine-specific TCRα clones from the antigen-specific TCR library
E) Longitudinal TCRα clonal trajectories grouped into ten clusters representing distinct patterns of clonal expansion and persistence. Trajectories were identified by hierarchical clustering of the top 1,500 clones from each participant across all time points. Colored boxes indicate trajectory clusters significantly enriched for Env- or Gag-specific TCRα clones

ND, non-detectable

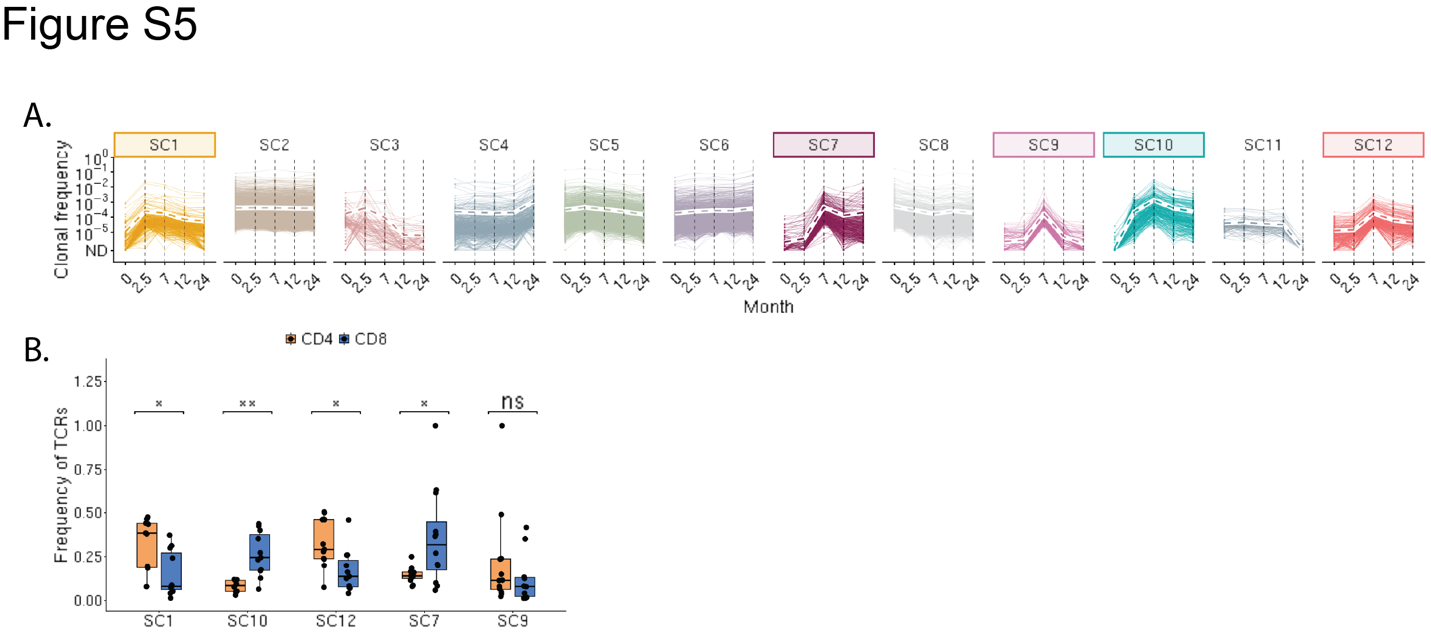

**Fig. S5. Longitudinal clonal trajectories and T cell subset composition of clones represented in the single-cell dataset**

A) Longitudinal vaccine-specific TCRβ and TCRα trajectory clusters identified by hierarchical clustering after restricting the longitudinal TCR dataset to clones represented in the single-cell dataset. Colored boxes indicate trajectory clusters significantly enriched for vaccine-specific clones
B) Participant-level proportions of unique TCR clones belonging to each CD4⁺ or CD8⁺ T cell population, as identified by single-cell profiling, within each trajectory cluster. Differences between trajectory clusters were assessed using Wilcoxon rank-sum tests with BH correction for multiple comparisons

*P < 0.05; ** P < 0.01; ns, not significant; ND, non-detectable

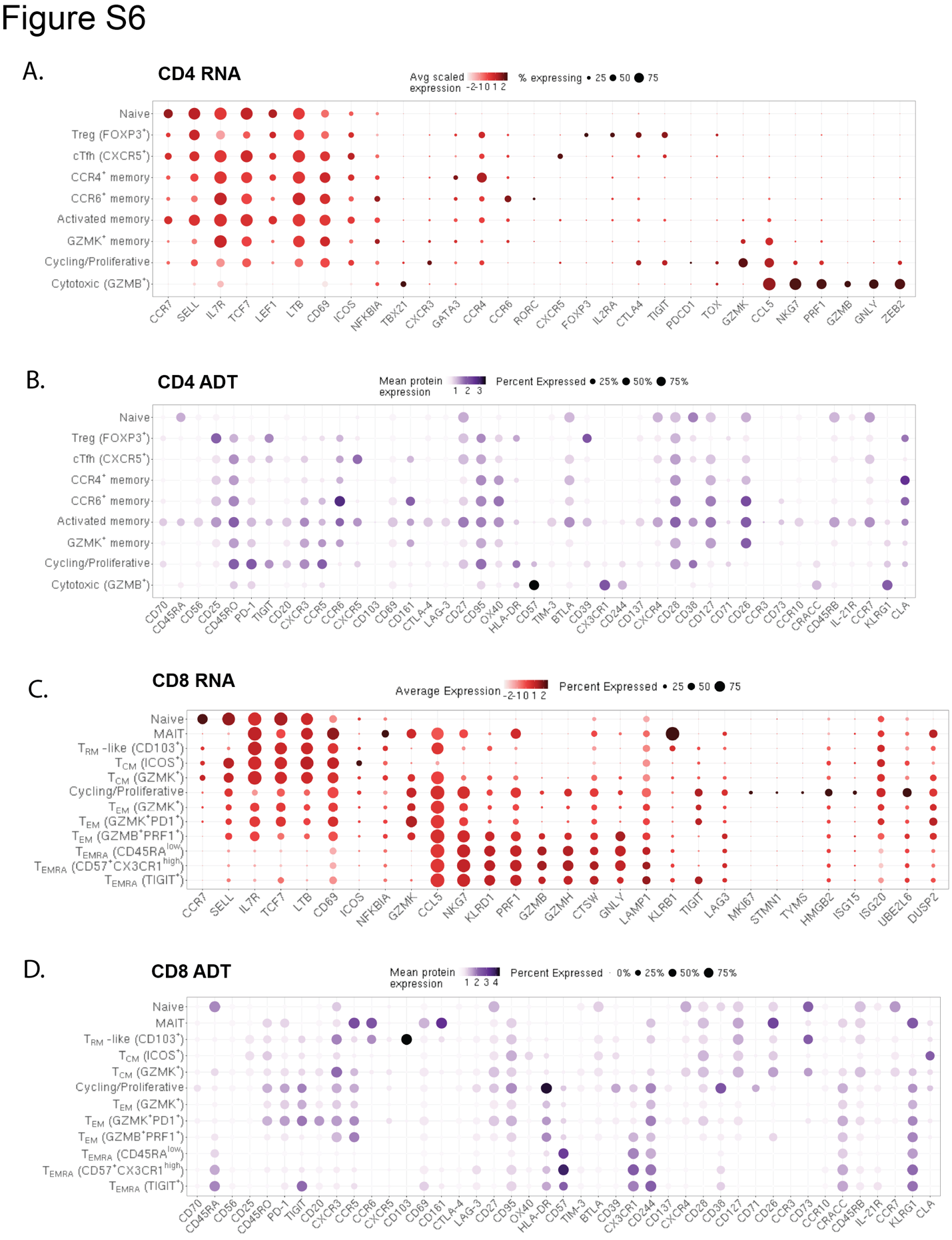

**Fig. S6. Validation of CD4⁺ and CD8⁺ T cell state annotations by transcriptional and surface protein profiles**

A) Dot plot showing RNA expression of canonical marker genes supporting CD4⁺ T cell state annotations. Dot size represents the percentage of cells expressing each gene, and color indicates average scaled RNA expression
B) Dot plot showing ADT expression profiles supporting CD4⁺ T cell state annotations. ADT marker positivity was defined using empirically determined expression thresholds (see Supplemental Materials and Methods). All measured surface protein markers were visualized across annotated CD4⁺ T cell states. Dot size represents the percentage of ADT-positive cells, and color indicates relative ADT expression across states
C) RNA expression of marker genes defining the annotated CD8⁺ T cell states. The expression profiles highlight transcriptional programs associated with the distinct CD8⁺ T cell states. Dot size represents the percentage of cells expressing each gene, and color indicates average scaled RNA expression
D) Surface protein expression profiles across annotated CD8⁺ T cell states. ADT marker positivity was defined using empirically determined expression thresholds (Supplemental Materials and Methods). Dot size represents the percentage of ADT-positive cells, and color indicates relative ADT expression across states

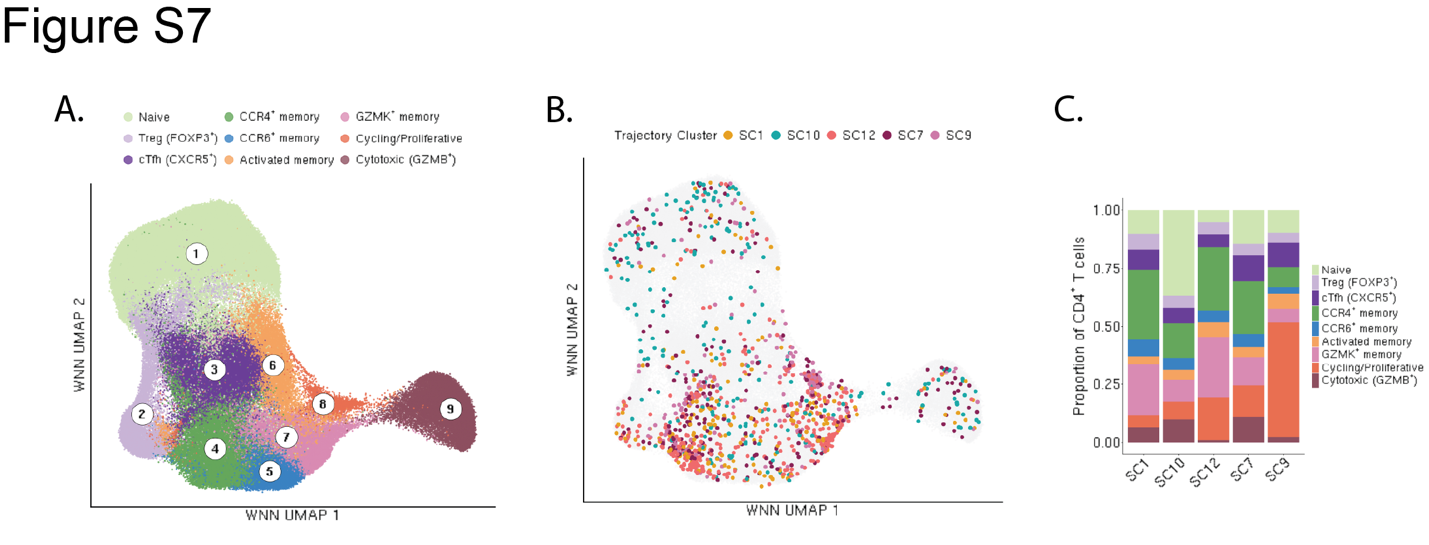

**Fig. S7. CD4⁺ T cell states across longitudinal clonal trajectories**

A) CD4⁺ T cell UMAP colored by annotated T cell states
B) CD4⁺ T cell UMAP colored by longitudinal trajectory clonal clusters
C) Stacked bar plot showing the proportion of annotated CD4⁺ T cell states within each longitudinal clonal trajectory cluster

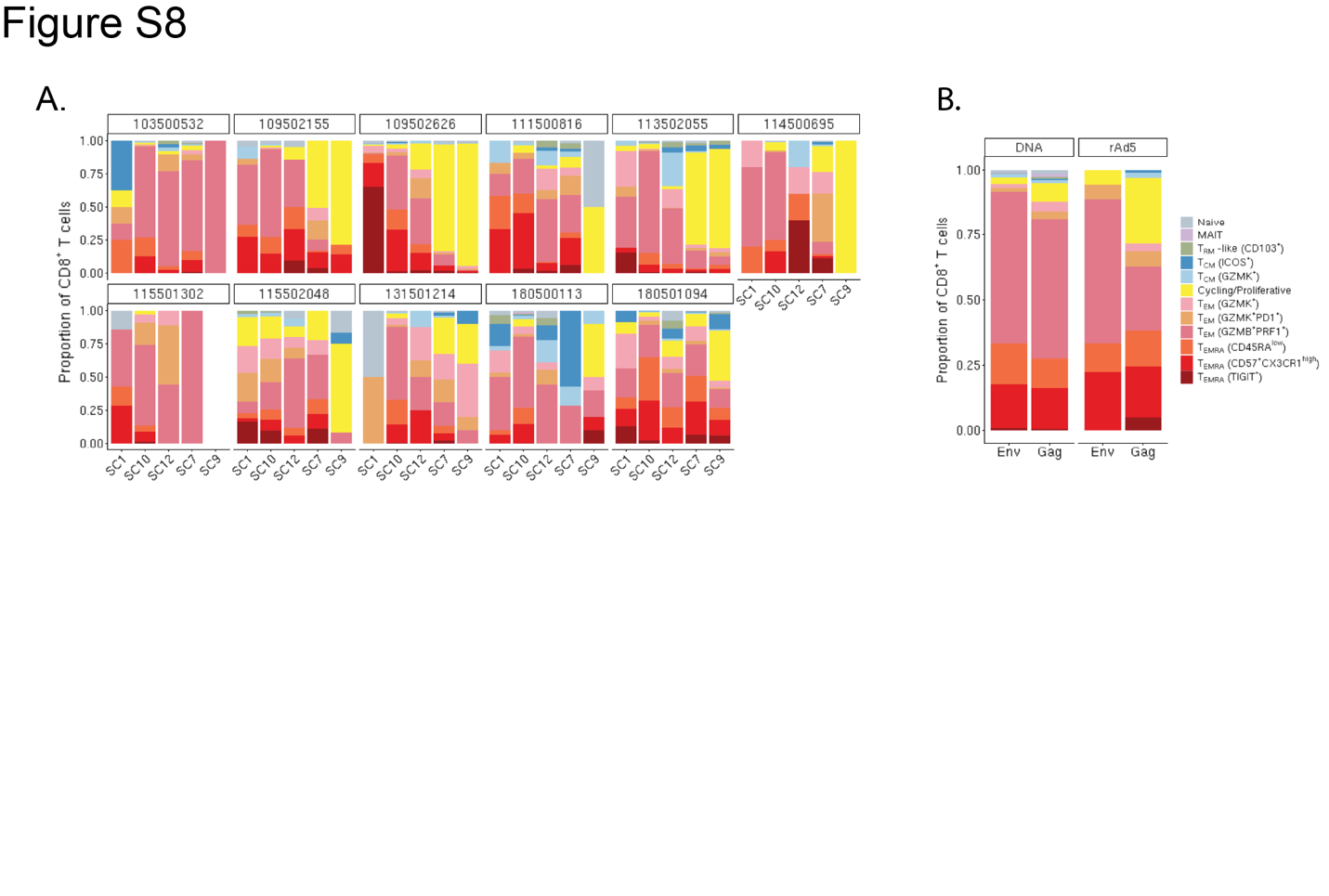

**Fig. S8. CD8⁺ T cell states associated with clonal trajectories**

A) Stacked bar plots showing the proportion of CD8⁺ T cell states within each longitudinal trajectory cluster for each participant
B) Proportion of CD8⁺ T cell states among DNA prime- and rAd5 boost-induced Env- and Gag-specific cells that remained detectable at M24
